# Temporal Patterns of Brain Network Plasticity During the Onset and Maintenance of Neuropathic Pain in Male Mice

**DOI:** 10.64898/2026.01.15.699728

**Authors:** Silvia Cazzanelli, Samuel Le Meur-Diebolt, Youenn Travert-Jouanneau, Luc Eglin, Ipek Yalcin, Adrien Bertolo, Alexandre Dizeux, Nathalie Ialy-Radio, Thomas Deffieux, Mickael Tanter, Bruno Felix Osmanski, Jeremy Ferrier, Sophie Pezet

## Abstract

Neuropathic pain arises from complex peripheral and central mechanisms and induces long-lasting maladaptive changes in the brain. To investigate the temporal dynamics of these changes, we examined resting-state functional connectivity (FC) in a mouse model of neuropathic pain across its initiation and maintenance phases.

Using functional ultrasound imaging to capture whole-brain FC over disease progression, we conducted two complementary studies: a longitudinal assessment in anesthetized animals and an analysis of awake cohorts at distinct disease stages. Both exploratory and literature-guided analyses revealed that FC across most large-scale networks remained remarkably stable during pain maintenance. In contrast, pain onset was marked by enhanced FC between key regions involved in sensory, emotional, and motivational processing, including the motor cortex and nucleus accumbens, the prelimbic and insular cortices, and the infralimbic cortex and hypothalamus. As pain persisted, we observed reduced FC within the somatomotor network, reflecting functional disconnection. Additionally, postsurgical pain alone produced enduring FC changes within the prefrontal cortex, hippocampus, and insula, indicating long-term central plasticity independent of neuropathic injury.

Together, these findings reveal dynamic, network-specific adaptations that distinguish the onset and maintenance phases of neuropathic pain and surgery-related plasticity.

## Introduction

Pain is defined as an unpleasant sensory and emotional experience associated with, or resembling that associated with, actual or potential tissue damage, and it often arises from a lesion or disease affecting the somatosensory system. Neuropathic pain represents a significant global health burden, affecting 7–10% of the population ^1^. In this condition, lesioned sensory neurons transmit abnormal signals to the spinal cord and brain, producing characteristic symptoms such as allodynia and hyperalgesia ^2^. Over time, this altered signaling induces long-lasting central sensitization, leading to widespread electrophysiological, structural, and functional changes across various brain regions ^3^.

These brain regions are often part of broader functional networks that serve multiple roles beyond pain perception, including emotional regulation and cognitive control. For instance, the cingulate cortex and the prefrontal-amygdala axis are critically involved in both nociception and mood regulation. This overlap has led to the hypothesis that maladaptive plasticity in these hubs may underlie the high comorbidity of chronic pain with affective disorders such as anxiety and depression ^4^.

While many studies have identified the brain regions activated during acute pain (often referred to as the ’pain matrix’), much less is known about how brain networks reorganize as pain becomes chronic. Pioneering work by Apkarian and colleagues revealed that functional connectivity (FC) between the nucleus accumbens (NAc) and other limbic and cortical structures could predict the transition to chronic pain in humans^5^, suggesting that network-level biomarkers may play a central role in pain chronification.

Resting-state functional connectivity has emerged as a promising approach for studying such brain-wide alterations. This method captures spontaneous fluctuations in neural activity, offering insights into the intrinsic organization of the brain and its longitudinal dynamics in both humans and animals ^6–8^. In preclinical models, it allows researchers to track neural adaptations in controlled environments and over defined disease stages.

Functional ultrasound (fUS) imaging is a powerful tool for this purpose, offering exceptional spatiotemporal resolution and the ability to image deep brain regions in both anesthetized and awake animals. Previous studies using fUS demonstrated its sensitivity to FC alterations in anesthetized rodents ^9–12^, as well as its feasibility in awake imaging conditions ^13–15^.

In this study, we applied fUS imaging to investigate the longitudinal evolution of resting-state FC in a mouse model of neuropathic pain ^16^, focusing on the initiation and chronification phases. By imaging both anesthetized and awake cohorts across multiple time points, we aimed to capture time-specific, network-level neuroplasticity associated with persistent pain. We first conducted an unbiased exploratory whole-brain analysis, followed by literature-informed secondary analyses.

Our results reveal time-specific selective changes in functional connectivity within brain networks involved in pain processing and emotional regulation, occurring during either the initiation or maintenance phases of neuropathic pain.

## MATERIALS AND METHODS

### Animals

The experiments were conducted in compliance with the European Community Council Directive of September 22nd 2010 (010/63/UE) and the local ethics committee (Comité d’éthique en matière d’expérimentation animale N° 59, “Paris Centre et Sud”, project # 19701 2019031020578789 V5). Accordingly, the number of animals in our study was kept to the minimum necessary. Based on previous studies using a similar experimental design ^15^, we established that N=6 animals per group was the minimum number of animals required to detect statistically significant differences in our imaging experiments. Using the intra and inter-animal variability established by Rabut et al., ^15^ and the variations in static functional connectivity measured in a previous study in awake freely moving mice ^14^, we performed a G-Power calculation (https://www.psychologie.hhu.de/arbeitsgruppen/allgemeine-psychologie-und-arbeitspsychologie/gpower). This analysis determined that sample sizes of N=6 to 8 animals per group were required. Finally, all methods were conducted in accordance with ARRIVE guidelines.

In total, 88 male C57BL/6 mice (Charles River Lab) were used, including 24 anesthetized and 64 awake animals.

Mice arrived in the laboratory one week before the beginning of the experiments at the age of 7 weeks and weighed between 20-25g. Animals were socially housed in well-ventilated cages. The housing room was kept at a constant temperature of 22°C with relative humidity kept between 45 and 50%. Food and water were provided *ad libitum*. The housing room was kept on a 12-hours light/dark reverse cycle (lights on from 8 pm to 8 am), and all experiments were conducted during the dark phase under red light.

### Sex as a biological variable

In this study, only male mice were included. This decision was based on two primary considerations: (i) the aim to minimize the number of animals used, in accordance with the 3Rs principles (Replacement, Reduction, Refinement) and current European regulations that place strict emphasis on reducing animal use; and (ii) the high complexity, duration, and resource intensity of the functional neuroimaging experiments, which spanned nearly four years of work.

To limit variability and focus on clearly interpretable effects, we prioritized one sex, acknowledging that male mice tend to exhibit lower inter-individual variability in the behavioral and neuroimaging phenotypes assessed.

We recognize that the exclusion of female animals limits the generalizability of our findings and that sex differences in neuropathic pain mechanisms are increasingly reported. Future studies will be necessary to determine whether the observed brain network alterations are conserved in female mice.

### Experimental designs

This study was conducted under two conditions: anesthetized and awake, resulting in two distinct experimental designs with corresponding timelines (Figure1).

#### Experiments performed in anesthetized animals (Figure 1A, 1C-E)

Anesthetized animals were randomly assigned to three groups. The “neuropathic” group (N = 8) underwent cuff surgery (Figure 1B) (explained in detail in the following paragraphs), consisting of implanting a polyethylene tube around the main branch of the sciatic nerve; The “sham” group (N = 8) underwent the same surgical protocol but did not receive the cuff implantation; The “naïve” group (N = 8) did not undergo any surgical procedures. Changes in functional connectivity was studied longitudinally before group assignment (T0) and at 2 weeks, 8 weeks, and 12 weeks post-surgery, which are different time points of neuropathic pain installation for the first one and chronification for the others (Figure 1A left).

**Figure 1:**
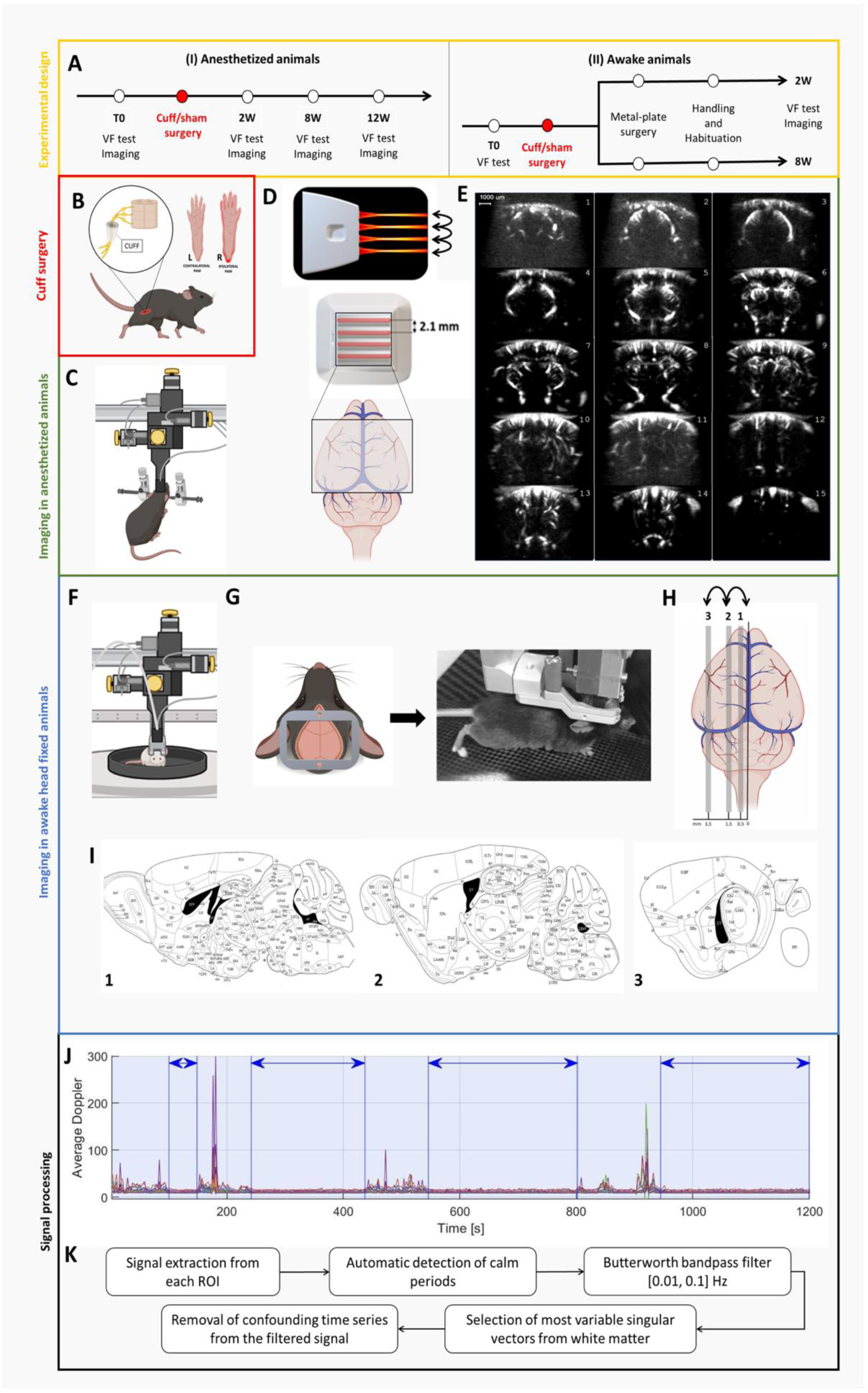
Experimental designs. A: Experimental design and projects timelines: (i) in anesthetized animals n=24 and (ii) in awake n=64, head-fixed animals. B: Schematic illustrating the animal model of neuropathic pain used in this study, involving the cuffing of the right sciatic nerve. C-E: Details of the experiments in anesthetized animals, including the motorized setup for precise probe displacement (C). D: Multi-array probe that includes four acoustic lenses, separated by 2.1 mm. This probe, along with its controlled motion, extends the field of imaging to encompass the entire cerebrum. E: Representative power Doppler images of the 16 imaging planes obtained. F-I: Details on the experiments in awake animals. F: Schematic representation of the imaging setup for the awake, head-fixed configuration. The animal is head-restrained while resting on a lightweight homecage that can move and rotate, providing a sensation of displacement, although the mouse remains fixed. The ultrasound linear probe is mounted on 3D motors allowing repetitive imaging in the three imaging planes of interest (H, I). I: Anatomical atlas depicting the three sagittal planes of interest. J: First motion artifacts are detected and removed. Only the clean signal is kept (blue arrows in the example). K: Main steps of the signal processing. (The panels B to F were created with Biorender.com).

At these four specific time points examined, the mechanical sensitivity of the hind paws was assessed on the day of imaging using the Von Frey test. Changes of resting-state functional connectivity were measured in the entire mouse cerebrum (Figure 1E, Bregma + 4 mm to Bregma -4 mm) of lightly anesthetized mice with a multi-array transducer ^17^ (Figure 1D).

During the course of this longitudinal study, several mice had to be excluded at various points along the project timeline (specifically: n = 2 at 2 weeks, n = 2 at 8 weeks, and n = 1 at 12 weeks). A detailed overview of the project timeline and the sequential phases of data collection and animal exclusion is provided in detail in Supplementary Figure 1.

#### Experiments performed in awake head fixed animals (Figure 1A, 1F-I)

In our preliminary experiments to establish functional connectivity (FC) imaging in awake animals, we initially aimed to perform longitudinal imaging in the same head-fixed mice for up to 12 weeks after nerve injury (14 weeks after metal plate implantation). However, after multiple trials and optimizations, we found this approach unfeasible. Although the imaging window remained intact for several weeks, repeated application of the imaging gel gradually softened the skull and eventually led to infection, despite careful cleaning and disinfection procedures. Consequently, we adopted an alternative strategy, imaging separate cohorts of animals in which the head plate was implanted two weeks prior to imaging. This approach consistently yielded high-quality imaging windows.

Awake animals were randomly assigned to three subgroups (naïve, sham and cuff). Their threshold of hind paw mechanical sensitivity was determined first at baseline (T0, i.e. before surgery) using the von Frey test (Figure 1A, right). Mice (N=64 in total) were then dispatched randomly in different groups: N=24 received sham surgery and N=26 underwent nerve cuffing (Figure 1B). The remaining 14 mice (naïve group) were not subjected to any surgical procedure. Three weeks before awake imaging, a metal plate was implanted on the skull of each mouse (Figure 1F, G), followed by habituation to restrained conditions. The study involved two separate cohorts examined at different time points: 2 weeks and 8 weeks post-surgery. At each time-point, the mechanical sensitivity of the hind paws was assessed on the day of imaging with the Von Frey test and changes in resting-state functional connectivity were evaluated in three alternating sagittal planes in awake head-fixed mice, using a motorized linear ultrasound probe (Figure 1H-I), following a well-established protocol ^18^. Each animal underwent multiple imaging sessions per time point, resulting in numerous acquisitions for each animal. For the analysis n=103 acquisitions were included, corresponding to n=54 for the 2W time point (naive n=15, sham n=18, NP n=21) and n=49 for the 8W time point (naive n=20, sham n=14, NP n=15).

During the study, several mice were excluded from the analysis (21 in the 2-week cohort and 22 in the 8-week cohort). Exclusions were due to one or more of the following reasons: (1) death of the animal, (2) outlier mechanical sensitivity thresholds, (3) aggressive behavior, (4) excessive motion leading to inadequate signal quality—only motion-free periods lasting at least 20 seconds were retained for analysis (see the section Power Doppler denoising below), and (5) poor-quality cranial window. Individual exclusion criteria are provided in Supplementary Figure 2.

### Measure of the threshold of mechanical sensitivity: Von Frey test

Mice were placed in Plexiglas boxes (9 cm x 7 cm x 7 cm) with a wire mesh floor. Following a 45-min habituation period, the mechanical threshold was assessed by measuring the withdrawal threshold of the stimulated paw in response to calibrated plastic filaments applied with increasing force (Von Frey filaments, from 0.16 g to 10 g). Each filament was tested five times per paw, applied until it bent (Yalcin et al. 2014; Barthas et al. 2015). The threshold was defined as three or more withdrawals observed out of the five trials.

### Surgical procedures

For all surgical procedures, a mixture of ketamine (100 mg/kg) and medetomidine (1 mg/kg) was administered intraperitoneally. As previously described ^17^, the eyes of the mouse were protected using an ointment (Ocry-gel, TVM, UK). Body temperature was controlled with a rectal probe connected to a heating pad set at 37 °C. Respiration and heart rate were monitored using a PowerLab data acquisition system with the LabChart software (ADInstruments, USA). The surgery duration ranged from 30 minutes for sciatic nerve cuffing and sham-operated animals to 45-60 minutes for metal plate fixation. At the end of the procedure, subcutaneous injections of atipamezole (1 mg/kg, Antisedan) and Metacam (10 mg/kg/day) were given to reverse the anesthesia and to prevent postsurgical pain, respectively, along with 0.1 ml of 2.5% Glucose. Animals were left in a warm chamber to recover from anesthesia. After recovery, they were returned to their cage. The wound healing process was monitored daily for one week.

#### Induction of the model of neuropathic pain (sciatic nerve cuffing)

As described in the early reports ^19^, once the animal was deeply anesthetized, it was positioned laterally on the surgical table to allow access to the right leg. The skin of the right leg was shaved and disinfected using Betadine®. A 0.5 cm incision was done in the skin and muscle of the leg, and a 2 mm-long PE-20 polyethylene tubing was placed around the main branch of the sciatic nerve of the right leg (Figure 1B). Animals in the sham group underwent the same procedure without cuff implantation. The skin was finally sutured using Prolene Ethicon 6-0 thread.

#### Implantation of metal plate for head-fixation

Once anesthetized, the mouse was placed on a stereotaxic frame and the skin was shaved, disinfected with Betadine®, and anesthetized locally with a subcutaneous injection of lidocaine (4 mg/kg). After a longitudinal incision from behind the occipital bone to the beginning of the nasal bone, the skull bone was exposed, and the periosteum was removed under aseptic conditions. The metal plate (sterilized using Alkacide instrument disinfectant) was fixed on the skull using Superbond C&B (Sun Medical, USA), Microhybrid composite (Henry Schein France) and small screws (Ref: #00-96X3-32, Phymep, France) minimally drilled into the skull ^18^. Once the metal plate was implanted, the skull bone was covered by a thin layer of surgical glue (Vetbond). The field of interest was approximately 5 mm wide, located between the Bregma and Lambda points. A protective cap was mounted on the metal plate using magnets to protect the skull and to keep the imaging field intact for several weeks ^18^. Altogether, the metal plate and the cap did not interfere with the normal daily activity of the mice. After a recovery period of one week, the mice proceeded towards the habituation phase.

### Fus imaging

#### Animal preparation for the imaging experiments in anesthetized mice

Induction of the anesthesia was performed by intraperitoneal injection of a mix of ketamine (80 mg/kg) and xylazine (8 mg/kg). As previously described elsewhere ^18^, its maintenance consisted in intramuscular perfusion of a mix of ketamine (15 mg/kg/h) and xylazine (1.5 mg/kg/h) via syringe pump, supplemented with 0.5% isoflurane (100% air as a carrying gas). The mouse’s hair was removed using depilatory cream, the skin was disinfected with Betadine®). The animal was positioned on a stereotaxic frame, and the scalp was incised along the sagittal suture, extending from the occipital bone to the nasal bone. Eyes were protected with an ophthalmic ointment (Ocry-gel, TVM, UK). Body temperature was monitored using a rectal probe connected to a heating pad set at 37°C. Respiration and heart rate were monitored using a PowerLab data acquisition system with LabChart software (ADInstruments, USA).

As we previously established that multiple depilation induces the growth of ingrown hairs and that they impact considerably the quality of images, after injection of a local anesthetic lidocaine (4 mg/kg), the skin was cut and sutured back at the end of the imaging session (using Prolene 6-0 polypropylene thread, Ethicon, USA).

#### Animal preparation for awake head fixed imaging experiments

##### Habituation to the head-restrained set-up

Mice were habituated to head restraint in the Mobile HomeCage (MHC, Neurotar, Finland), which combines stable head fixation with a carbon cage suspended on an air cushion, allowing free movement during restraint. This setup provides a familiar, low-stress environment for the animals. Training included two phases: (1) daily handling (10–15 min) for at least four days starting two days post-surgery, followed by 5 minutes of free exploration in the MHC, and (2) progressive head-fixation habituation over at least five days. Head-fixation duration increased daily from 15 minutes to 2 hours, with total training duration adjusted per animal. Most mice were considered ready for imaging after 5 days of habituation. However, some required an additional two days of habituation, as previously observed ^18^. Mice were weighed daily, with >20% weight loss as an exclusion criterion—not observed in this study. Positive reinforcement (Honey Pops, Kellogg’s) was provided after each session to reduce stress. Background music was played continuously to mask setup noise and further minimize stress.

#### fUS imaging

Following preparation, the skull of the animal was gently covered with centrifuged sterile gel to facilitate acoustic coupling with the fUS probe.

The acquisitions were performed using a prototype ultrasonic ultrafast neuroimaging system (Iconeus, Paris, France - ART Inserm Ultrasons Biomédicaux). A coronal scan of the entire window (Bregma to Lambda) was first performed to reconstruct a 3D angiographic volume of the brain. Using the IcoStudio software (Iconeus, Paris, France), we automatically aligned each individual 3D angiography with the Allen Mouse Brain Common Coordinate Framework ^20^ ensuring consistent fUS volume acquisition across animals and imaging sessions.

Functional connectivity was assessed at an adjusted sampling rate of 0.55 Hz, enabled by repeated plane acquisitions using a motorized positioning system and vascular pattern recognition.. The detailed procedure has been previously described in ^18^.

#### Choice of the imaging fields

In both sets of experiments, imaging sessions consisted of 20-min acquisitions, either in the coronal planes (anesthetized animals) or parasagittal planes (awake head fixed animals). FUS imaging currently lacks simultaneous imaging in three spatial dimensions. Nevertheless, by combining these two experimental configurations, this study explored the changes of FC within a pseudo-3D volume.

The difference in imaging planes between the two experimental designs reflects a technical limitation at the time the awake imaging experiments were initiated in 2021, when multi-array probes were not yet available. Under these conditions, imaging in the sagittal plane represented the most effective strategy to capture FC across the regions of interest in awake animals.

#### Imaging field in anesthetized configuration

We first aimed to monitor changes in FC across a large part of the mouse’s cerebrum in a longitudinal setting. For this purpose, we used a recently developed probe (IcoPrime 4D-MultiArray, Iconeus, Paris, France) which features four compact 64-element linear arrays (110 μm pitch) spaced 2.1 mm apart to reduce crosstalk and optimize coverage ^17^. Mounted on a 4-axis motorized stage, the probe sequentially scanned the brain in 0.525 mm steps along the antero-posterior axis, acquiring a full volume of 10 x 8.1 x 9 mm^3^ comprised of 16 Power Doppler images of 110 x 100 x 525 µm^3 resolution, as described previously ^17^. After slice timing correction, each full mouse brain volume was acquired in 2.4 s (Figure 1E). The 40 ROI studied are shown in Supplementary Figure 3 and listed in Table 1A.

**Table 1:**
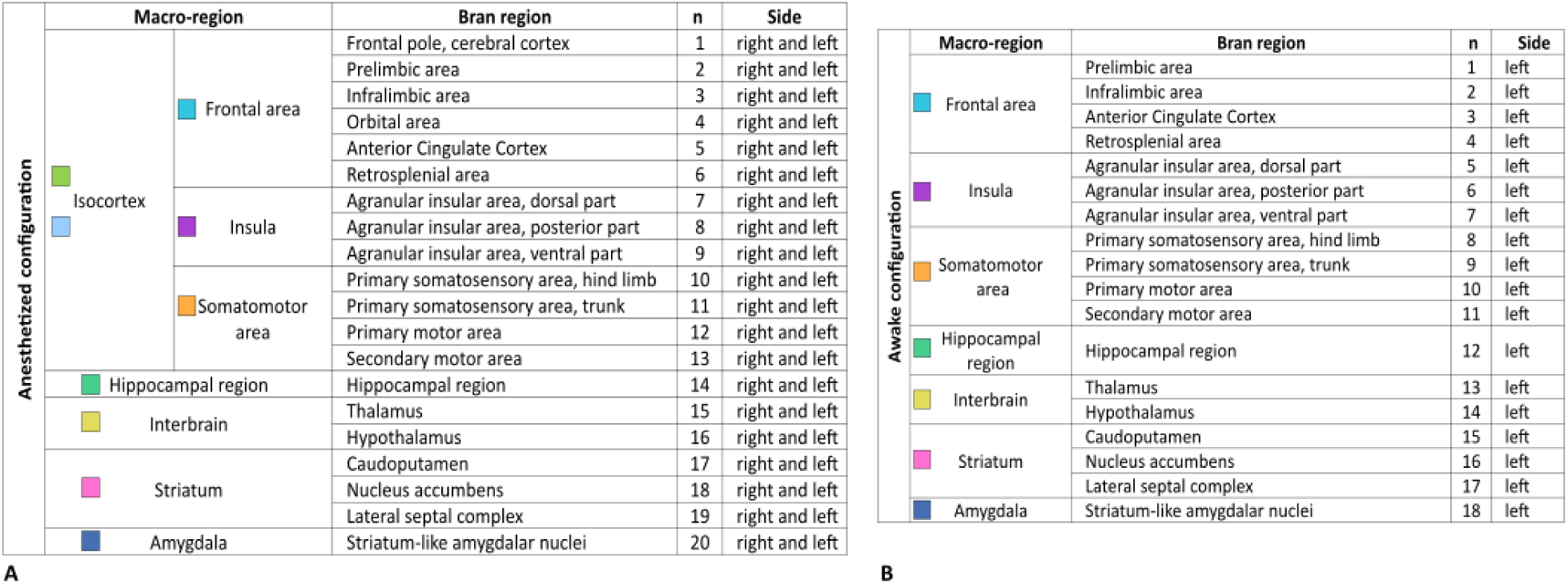
List of Regions of Interest (ROIs). (A) For the anesthetized configuration, the 40 ROIs are grouped into five macro-regions: the Isocortex, which includes the frontal area, insula, and somatomotor area; the hippocampal region; the interbrain; the striatum; and the amygdala. In this configuration, the brain was imaged bilaterally, with each ROI divided into right and left hemispheric parts. (B) For the awake configuration, the 18 ROIs are grouped into seven macro-regions: the frontal area, insula, somatomotor area, the hippocampal region, the interbrain, the striatum, and the amygdala. In this configuration, the brain was imaged on its left hemisphere.

#### Imaging field in awake configuration

The purpose of this study was to follow up the changes of FC in the left part of the mouse’s cerebrum, in awake conditions in a sagittal configuration. A 15 MHz linear probe was used to match this purpose. The probe was mounted on a 4-axis linear motor stage that stepped the probe sequentially along the sagittal axis (Figure 1H) within a 0.2 sec duration, allowing the acquisitions in three specific planes located at L=0.5 mm, L=1.5mm and L=3.5mm (Figure 1I). Based on the literature, we chose to focus our study on 18 specific brain regions (Supplementary Figure 3, Table 1B). Due to the expected possibility of signal loss due to motion artifacts, each animal underwent four to six 20-min runs during each imaging session.

#### fUS imaging sequences

The fUS imaging sequence consisted of eleven successive tilted ultrasonic plane waves with an angle ranging from −10° to 10° emitted at a 5.5 kHz Pulse Repetition Frequency (PRF) for the experiments in awake configuration and 4 × 8 tilted plane waves acquired at a pulse repetition frequency of 4 kHz (-12°, -8.57°, -5.14°, 1.71°, 1.71°, 5.14°, 8.57°, 12°) for the anesthetized configuration. For both parts of the study, the backscattered echoes were recorded by the transducer array and beamformed to produce blocks of 200 consecutive ultrafast images with a framerate of 500 Hz.

##### Signal processing

In order to filter the Cerebral Blood Volume (CBV) and to remove the tissue signal, we used a clutter filter based on Singular Value Decomposition (SVD) applied to 200 successive frames (that is, 400ms). We regressed out the singular vectors of highest energy, removing either 60 singular vectors for anesthetized experiments or 35 singular vectors for awake experiments. Singular vectors with high energy were previously found to correspond to clutter signals ^21^. Finally, a Power Doppler image was obtained by integrating the energy of the filtered frames. Taking into account the duration of the probe movements between planes, this resulted in a sampling frequency of 2.1 s for awake acquisitions and 2.4 s for anesthetized acquisitions.

#### Power Doppler denoising

Prior to statistical analysis, artifacts in the Doppler signal caused by background noise or animal movement were eliminated using dedicated signal processing, as shown in Figure 1 J-K. Power Doppler signals were denoised using the IcoLab software (ICONEUS, Paris) to mitigate the effects of motion artifacts (in awake acquisitions) and hemodynamic variations from non-neuronal sources. The denoising process involved the following steps:

1. Computation of confounding time series through principal component analysis (PCA) of the most variable time series obtained from white matter voxels, following the method outlined by ^22^. Specifically, the top 5% most variable voxels were selected, and their top 5 principal components were retained as confounding variables.
2. Detection of high motion volumes in awake acquisitions by identifying volumes where the smoothed global signal (i.e., the average power Doppler signal from brain voxels, smoothed using a 5-sample moving average) deviated by more than 5% from its baseline. The global signal baseline was determined using the least trimmed square regression. Only periods of low motion lasting at least 20 seconds were retained, with any remaining volumes interpolated to prevent bias in subsequent temporal filtering.
3. Temporal filtering of power Doppler signals and confounding time series using a Butterworth forward-backward bandpass filter with a frequency range of [0.01, 0.1] Hz and an order of 8.
4. Regression of filtered confounding time series from the power Doppler signals to remove confounding effects (Figure 1J).

#### Seed-based maps

As recommended ^23^, to verify the strength of FC, we verified the strength of functional connectivity in the somato-motor network using ICA analysis (Supplementary Figure 5). The analysis was performed through the software Icolab, using processing previously described by our group ^17^.

#### Correlation matrix analysis and statistical approach

As the aim of this study was to investigate the changes of FC in an unsupervised approach, we further studied changes of FC in regions of interest (ROI) defined by the Allen Mouse Brain Atlas. We used the brain positioning system described in Nouhoum et al., 2021 [40], implemented in the IcoStudio software (Iconeus, Paris, France). We previously demonstrated that this approach enhances reproducibility in probe placement and accuracy in the ROI delineation. In short, as described elsewhere ^24^, a linear scan (6 mm span along the anterior-posterior axis, 0.2 mm step size) was conducted at the beginning of each session, co-registered to an average scan, and aligned with a standard Doppler reference template pre--aligned to the Allen Mouse Brain Atlas common coordinate framework ^20^.

The resulting ROI signals were normalized, slice-timing-corrected, and denoised. Subject-level FC matrices were then computed using Pearson correlation of every ROI signal pair.

Finally, group-level FC matrices were computed by averaging Fisher-transformed subject-level matrices, followed by back-transformation to correlation coefficients, as previously described ^12,17^. If the transformed data were normally distributed, a parametric Welch test was performed; otherwise, the Mann-Whitney test was applied. We performed Benjamini-Hochberg’s correction for multiple comparisons, with a false discovery rate of 0.05. Unfortunately, no significant changes were observed using this unsupervised approach (Figures 2 and 4).

**Figure 2:**
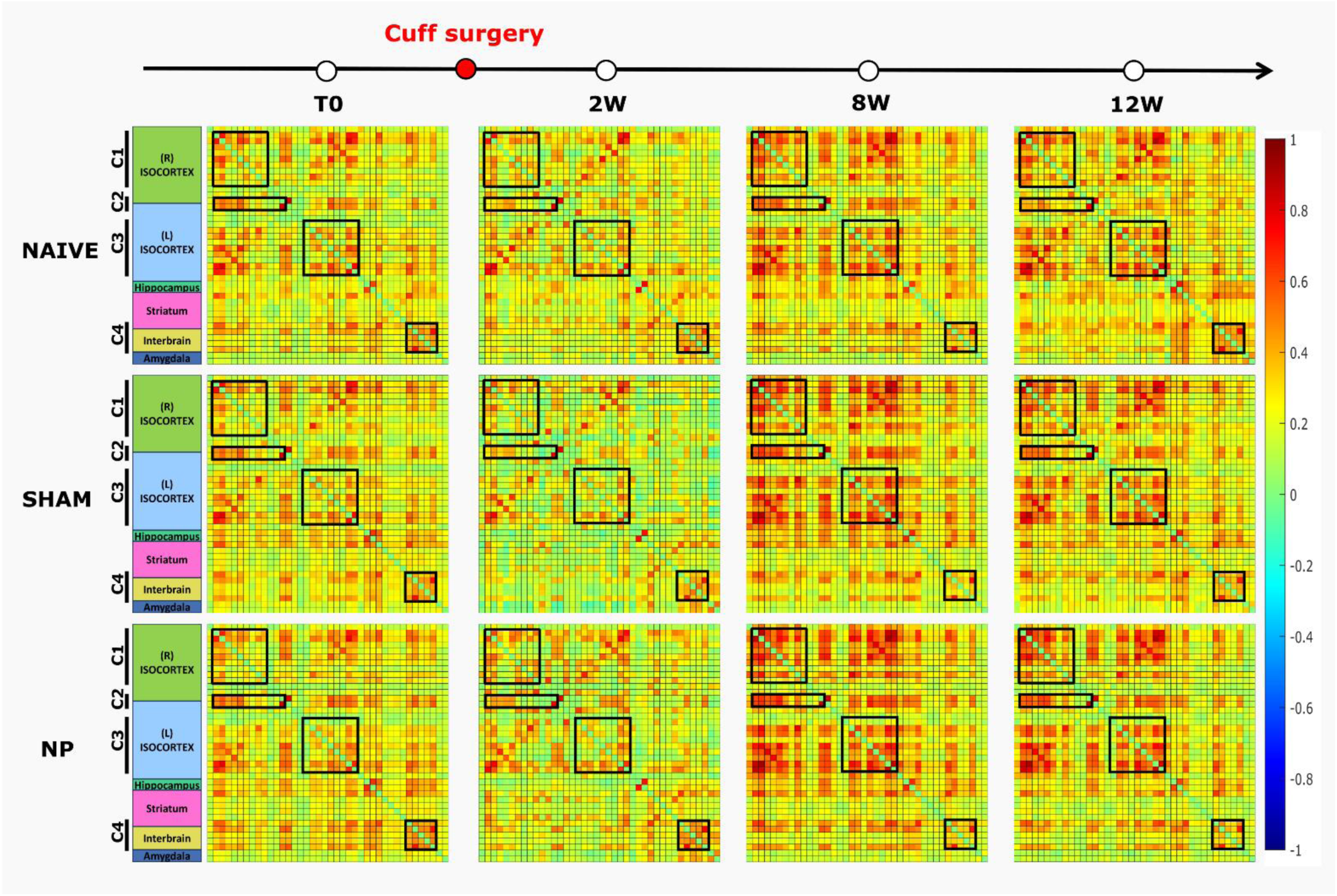
Functional connectivity alterations of wide-range networks in anesthetized mice. Group averaged resting-state functional connectivity matrices, displaying the mean Pearson correlation coefficients between 40 predefined regions of interest (ROIs; listed in Table 1A), computed for each experimental group (NAIVE, SHAM, NP) across four timepoints: baseline (T0), and 2, 8, and 12 weeks (2W, 8W, 12W) following cuff surgery. Color scale represents correlation values from -1 (blue, strong negative correlation) to +1 (red, strong positive correlation). Each matrix is organized by anatomical subdivisions (e.g., Isocortex, Hippocampus, etc.), and four network clusters (C1–C4) are highlighted by black squares. T0: Naïve n=8, Sham n=8, NP n=8; 2W: Naïve n=7, Sham n=7, NP n=8; 8W: Naïve n=8, Sham n=6, NP n=6; 12W: Naïve n=7, Sham n=6, NP n=6.

Alternatively, a second hypothesis-driven statistical approach was employed to assess potential changes in FC between selected pairs of ROIs across time points in the neuropathic and control groups. A two-way ANOVA (mixed model) was performed using GraphPad Prism 10. If a significant effect of treatment, time, or treatment-by-time interaction was detected (p < 0.05), post-hoc Tukey’s test with multiple comparisons correction was applied.

## RESULTS

### Development of mechanical allodynia

It was previously demonstrated that nerve injury caused by the cuff implantation results in mechanical hypersensitivity in the lesioned paw of the neuropathic group ^16^. At T0, no difference in mechanical sensitivity threshold was observed between groups, both in the anesthetized (Supplementary Figure 4A-B) and the awake conditions (Supplementary Figure 4C-D).

#### Anesthetized study

Although there was no difference in the mechanical threshold between groups before surgery, unilateral cuff implantation produced an ipsilateral mechanical allodynia that persisted for 12 weeks. The development of the unilateral allodynia is specific to the cuff implantation as we observed a significant decrease in the mechanical sensitivity threshold exclusively in the neuropathic group compared to the control groups (2W p=0.0002, 8W p=0.0021, 12W p=0.003) (Supplementary Figure 4A). On the other hand, no differences were observed in the mechanical sensitivity of the contralateral paw between the groups (Supplementary Figure 4B).

#### Awake study

Although there was no difference in the mechanical threshold between groups before surgery, unilateral cuff implantation produced an ipsilateral mechanical allodynia at 2W (Supplementary Figure 4C) and at 8W (Supplementary Figure 4E). We observed a significant decrease in the mechanical thresholds of the neuropathic group’s ipsilateral paw compared to the control groups’ ipsilateral paws (ipsiNAIVE/ipsiSHAM vs ipsiNP p<0.0001, Supplementary Figure 4C) (ipsiNAIVE/ipsiSHAM vs ipsiNP p= 0.0089, Supplementary Figure 4E). There is a significant difference between the threshold assessed in the ipsilateral paw within the neuropathic group before and after the surgery (p<0.0001, Supplementary Figure 4C) (p=0.0427, Supplementary Figure 4E). We also observed a significant decrease in the mechanical sensitivity threshold between contralateral and ipsilateral paw within the neuropathic group at both 2 weeks (contraNP vs ipsiNP p<0.0001, Supplementary Figure 4C) and 8 weeks (contraNP vs ipsiNP p=0.0015, Supplementary Figure 4E). On the other hand, no differences were observed in the mechanical sensitivity of the contralateral paw between the groups (Supplementary Figure 4D and 4F).

These results indicate the presence of mechanical allodynia in the lesioned paw of the neuropathic group, beginning at 2 weeks and persisting consistently at 8 and 12 weeks. The assessment of unilateral mechanical allodynia suggests that the neuropathic group exhibited typical neuropathic pain symptoms during the imaging sessions for both anesthetized and awake experimental conditions.

### FC alterations in a wide-range network in anesthetized mice

The group averaged FC, presented as correlation matrices in Figure 2 and as circular networks in Supplementary Figure 6, respectively, show a consistent and reproducible pattern of correlations across groups and timepoints, with trends of changes over time.

As previously described by several teams, a dominant feature of these connectivity patterns is a prominent cluster of strong inter-hemispheric correlations within the isocortex. This cluster further subdivides into three distinct subclusters (labeled as C1-2-3 in Figure2): somatomotor regions in the right hemisphere (C1), bilateral regions of the retrosplenial cortex (C2), somatomotor regions in the left hemisphere (C3). At baseline (T0), these subclusters showed strong connectivity across all groups. A marked reduction in correlation strength was observed at 2 weeks post-surgery (2W), followed by a pronounced increase at 8 and 12 weeks (8W and 12W), suggesting a dynamic reorganization of cortical connectivity over time in all groups of animals.

Additionally, robust intra-regional connectivity was observed, within the interbrain areas, across all conditions (labeled as C4 in Figure2). However, this connectivity slightly diminished at 2W, consistently across all groups, before returning to higher levels at later time points.

These findings underscore the reproducibility and reliability of the connectivity measurements and reveal specific clusters of temporal and group-dependent variability.

Visual inspection of the group-level connectivity, especially by focusing on the Sham vs NP and Naïve vs Sham group comparisons across timepoints, reveals distinct patterns of functional connectivity disruption. At baseline (T0), all groups display nearly identical FC profiles, indicating no pre-existing differences. At 2W: NP group shows an increased connectivity compared to sham and naive groups, especially in the retrosplenial cortex (C2), interbrain regions (C4), and bilateral isocortical subclusters (C1 and C3). At 8 and 12W: NP group connectivity is still stronger in some areas, especially between isocortical hemispheres (C2).

These visual inspections emphasize that the NP group shows greater and longer-lasting disruption of FC. The effects were regionally specific, with prominent differences in inter-hemispheric isocortex (C1 and C3), retrosplenial cortex (C2), and interbrain regions (C4). Differences between NP and Sham persisted longer (until at least 12W), unlike the transient changes seen between Sham and Naïve.

To statistically evaluate group-level differences underlying the observed connectivity patterns, we applied an unsupervised approach to assess changes in functional connectivity (FC) across the large-scale brain network (see *Materials and Methods*, section "Correlation Matrix Analysis and Statistical Approach"). However, these differences did not survive statistical testing after correction for multiple comparisons. We therefore report them as exploratory, descriptive trends only, not as evidence of confirmed group differences..

### Functional connectivity alterations in a specific subnetwork in anesthetized mice

Alternatively, a literature-informed secondary analysis was employed to assess potential changes in functional connectivity (FC) between selected pairs of regions of interest (ROIs) across time points in the neuropathic and control groups. This analysis revealed significant alterations in FC between several key regions, including the somatomotor (M1) and the striatum (NAc), the frontal area (IL), as well as the amygdala (Amy), the frontal area (PL) and the insular cortex (AIV) (Figure 3A).

**Figure 3:**
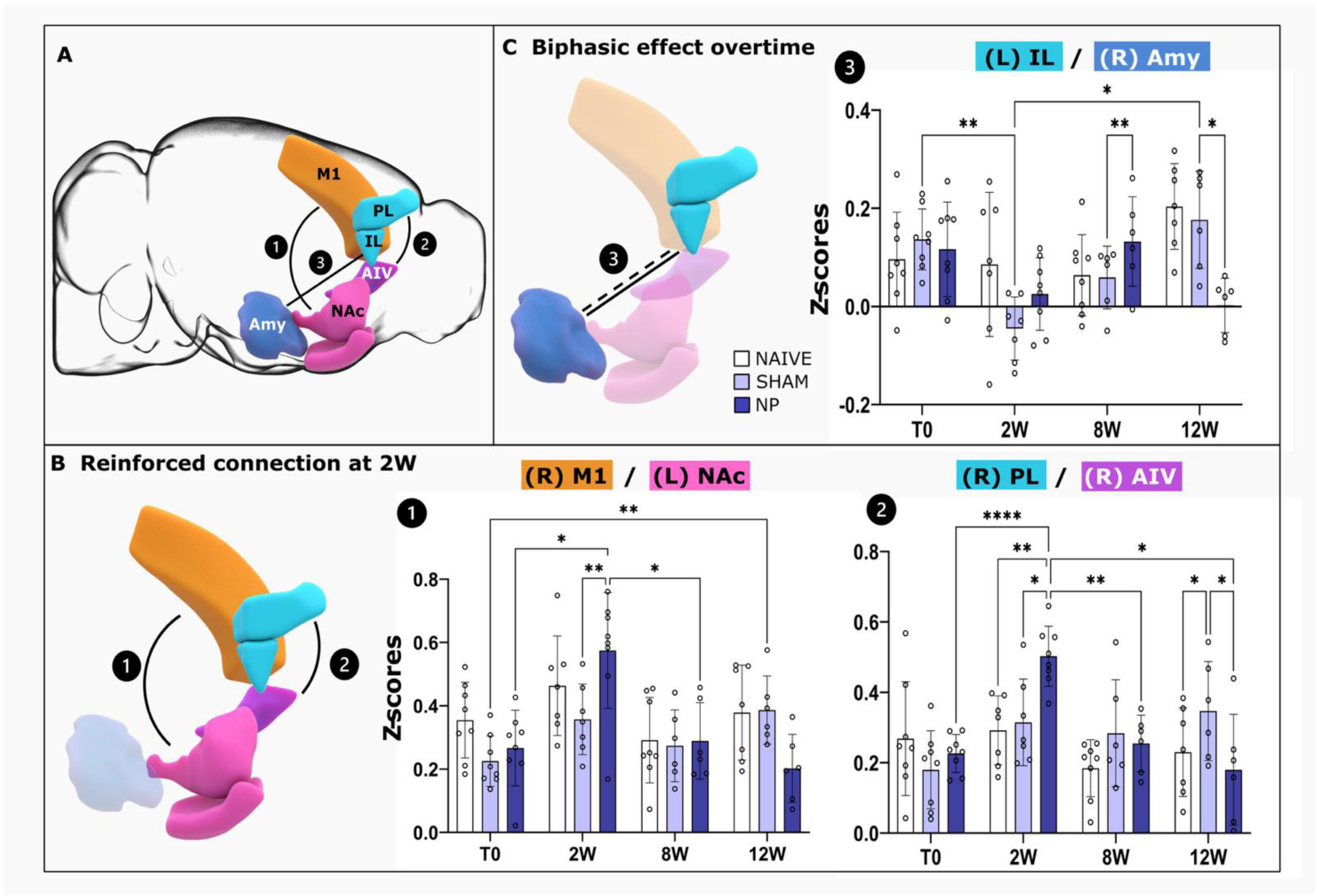
Functional connectivity alterations of specific subnetwork in anesthetized mice. (A) Schematized illustration of two subnetworks showing significant functional connectivity alterations: somatomotor area and striatum (1), and frontal area, amygdala and insular cortex (2-3). (B-C) Division of subnetwork showing a a variating effect over time (C). The increase in connectivity is illustrated as a bold line and the decrease as a dotted line. The changes illustrated refer to the alterations detected in the neuropathic group compared to the sham group. Graphs 1, 2 and 3 represent Fisher-transformed z-score of correlations between pairs of ROIs at four time points (T0, 2, 8 and 12 weeks). Data are presented as mean ± SD. *p<0.05, **p<0.01, ****p<0.0001. T0: Naïve n=8, Sham n=8, NP n=8; 2W: Naïve n=7, Sham n=7, NP n=8; 8W: Naïve n=8, Sham n=6, NP n=6; 12W: Naïve n=7, Sham n=6, NP n=6.

Specifically, connectivity between the right primary motor cortex (R-M1) and the left nucleus accumbens (L-NAc) (Figure 3B) was significantly increased at the onset of neuropathic pain (2W post-surgery), compared to both the sham group (p=0.0059, q=7.061, DF=6) and to the same animal baseline (T0 p=0.0151, q=6.039, DF=7). However, this enhanced connectivity did not persist during maintenance phase of chronic pain (Figure 3B, graph 1).

A similar transient increase in FC was observed between additional brain regions at the initiation of neuropathic pain. Specifically, connectivity between the right prelimbic cortex (R-PL) and the right anterior insular cortex (R-AIV) was significantly strengthened at this time point (Figure 3B, graph 2), with statistical comparisons showing increased FC relative to the sham group (p = 0.0209, q= 5.376, DF=6), the naïve group (p = 0.0056, q=7.152, DF=6), and the same animals at baseline (T0; p < 0.0001).

Secondly, certain networks exhibited variations over time of the FC changes between the left infralimbic cortex (L-IL) and the right amygdala (R-Amy) (Figure 3C).

During the maintenance phase of neuropathic pain (8- and 12-weeks post-injury), animals in the NP group displayed biphasic alterations in this emotion-related network. FC between the L-IL and R-Amy initially increased at 8 weeks compared to the SHAM group (p = 0.0087, q=8.449, DF=4), followed by a significant reduction at 12 weeks (p = 0.0344, q=5.669, DF=4 vs. SHAM; Figure 3C). These findings highlight distinct, time-dependent alterations in FC between neuropathic and control groups. Notably, we identified a specific pattern characterized by an early increase in connectivity between frontal regions and the amygdala during the maintenance of neuropathic pain (8 weeks), followed by a decrease in these connections during the chronification phase.

### Study of FC alterations in a wide-range network in awake mice

In order to prevent the potential interfering effect of anesthesia on FC, the second part of our study used awake, head restrained animals. In these experiments, many acquisitions were discarded due to strong motion artifacts (see Supplementary Figure 2). Using all acquisitions kept, the average connectivity networks are presented in Figure 4, as a correlation matrix and in Supplementary Figure 7, as circular networks.

**Figure 4:**
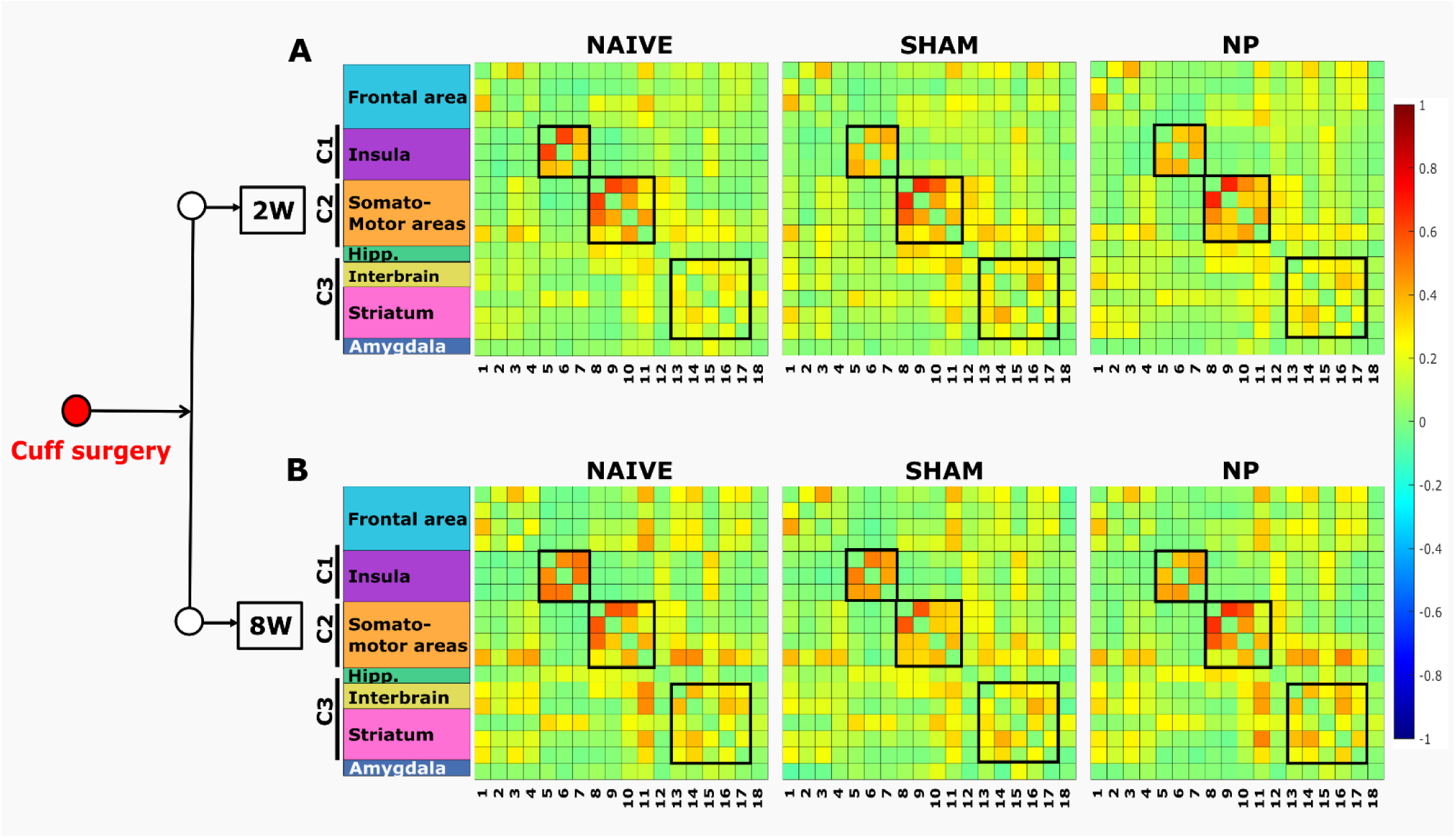
Functional connectivity alterations of wide-range networks in awake mice. (A-B) Group averaged resting-state correlation matrices, displaying the mean Pearson correlation coefficient between 18 selected ROIs (listed in Table 1B). Color scale represents correlation values from -1 (blue, strong negative correlation) to +1 (red, strong positive correlation). Each matrix is organized by anatomical subdivisions (e.g., Frontal area, Insula, etc.), and three network clusters (C1–C3) are highlighted by black squares. 2W: Naïve n=2, Sham n=5, NP n=5; 8W: Naïve n=5, Sham n=5, NP n=2.

In all groups, the connectivity pattern consistently revealed three distinct clusters (labeled as C1-2-3 in Figure 4): among the subregions of the insular cortex (C1), the somatomotor area (C2), and a striatum–interbrain (C3) cluster. Despite the shared network architecture, the strength of correlations within these clusters varied between groups at both 2W and 8W.

Visual inspection of the group-level connectivity reveals at 2 weeks (2W) in the insular cluster (C1), a strong correlation between its three subdivisions. The dorsal–posterior insula in the Naïve group shows the strongest internal connectivity. In contrast, both Sham and NP exhibited reduced connections between these subdivisions. In the somatomotor cluster (C2), the connectivity between the primary somatosensory area, trunk (S1Tr) and primary motor area (M1) with primary somatosensory area; hind limb (S1HL) is strong in both Naïve and Sham but appears less tightly connected in the NP group. However, the connection between S1Tr and secondary motor area (M2) is stronger in NP compared to Sham and Naïve. Regarding the striatum–interbrain cluster (C3), Sham and NP maintained a similar organization compared with Naïve, but with slightly stronger correlations (Figure 4 and Table1B).

At 8 weeks (8W) in the insular cluster (C1), the Naïve group continued to show stronger internal correlations compared to Sham and NP, though the intensity appears slightly increased relative to 2W. In the somatomotor cluster (C2), the connectivity between S1HL, S1Tr, and M1 was stronger in the NP group than in Sham or Naïve. In the striatum–interbrain cluster, NP showed enhanced correlations compared to Sham and Naïve (Figure 4 and Table1B).

These results demonstrate a consistent pattern of connectivity, indicating robust reproducibility across groups and time points. The robustness of this pattern enables us to identify specific clusters of variability in the strength of connections across groups.

To statistically evaluate group-level differences underlying the observed connectivity patterns, we applied an unsupervised approach to assess changes in functional connectivity (FC) across the large-scale brain network (see *Materials and Methods*, section "Correlation Matrix Analysis and Statistical Approach"). However, these differences did not survive statistical testing after correction for multiple comparisons. We therefore report them as exploratory, descriptive trends only, not as evidence of confirmed group differences.

### Functional connectivity alterations in specific subnetworks at early stage of neuropathic pain in awake mice

Alternatively, a literature-informed secondary analysis was employed to evaluate changes in functional connectivity (FC) between selected pairs of regions of interest (ROIs) across time points in neuropathic and control groups.

Among the FC alterations observed during the early phase of neuropathic pain, a significant increase in connectivity between the infralimbic cortex (IL) and the hypothalamus (Hyp, Figure 5A) was detected specifically in neuropathic animals (p=0.0069, q=4.396, DF=97 vs naive) but not in sham animals (p=0.0152, q=4.013, DF=97 vs sham), suggesting a neuropathic pain-specific pattern of brain plasticity (Figure 5B).

**Figure 5:**
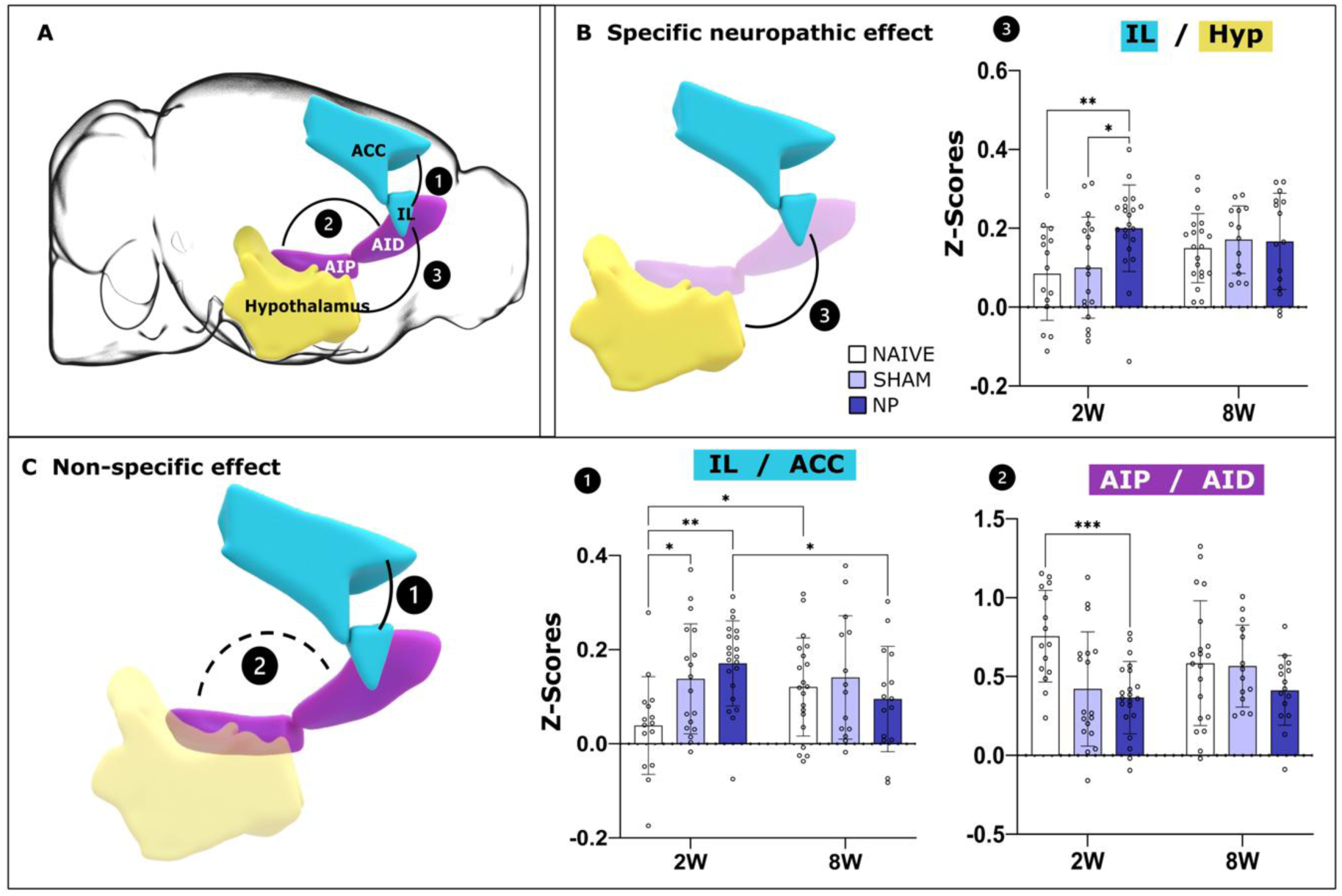
Functional connectivity alterations of specific subnetworks at 2W post-lesion in awake neuropathic mice. (A) Schematized illustration of the two subnetworks showing significant functional connectivity alterations. Frontal area, the interbrain and the interconnections between the posterior and the dorsal part of the insular cortex. (B-C) Division of subnetwork showing a specific neuropathic effect (B) and a non-specific effect (C). The increase in connectivity is illustrated as a bold line and the decrease as a dotted line. The changes illustrated refer to the alterations detected in the neuropathic group compared to the sham group. Graphs 1, 2 and 3 represent Fisher-transformed z-score of correlations between pairs of ROIs at 2 and 8 weeks. Data are presented as mean ± SD (*p<0.05, **p<0.01, ***p<0.001). Mice n=12, acquisitions n=54. 2W: Naïve n=2, Sham n=5, NP n=5.

In contrast, FC alterations common to both sham and neuropathic groups were identified across two pairs of regions involving the frontal cortex (IL and anterior cingulate cortex, ACC), and the insular cortex (AIP and AID, Figure 5C). These shared alterations suggest changes related to post-surgical pain rather than neuropathic pain specifically.

The specific pairs of regions showing these common FC alterations are as follows: 1) Infralimbic cortex (IL) and anterior cingulate cortex (ACC): FC was significantly altered in both the sham and NP groups compared to naïve animals (p = 0.0314, q=3.676, DF=57 naïve vs. sham; p = 0.002, q=5.066, DF=57 naïve vs. NP; Figure 5C, graph 1). 2) Two subregions of the insular cortex (AIP and AID): Connectivity between these regions was reduced in both sham (p=0.0062, q=4.447, DF=97) and NP (p = 0.0008, q=5.352, DF=97) animals compared to naïve controls, with no significant differences between the sham and NP groups (Figure 5C, graph 3).

These findings indicate that some of the observed FC changes may reflect non-specific effects related to surgery and post-surgical pain rather than neuropathic pain itself at early stage of cuff surgery.

### Functional connectivity alterations in specific subnetworks during persistent neuropathic pain in awake mice

During neuropathic pain maintenance (eight weeks after the cuff surgery) a distinct subnetwork of significant connectivity alterations was identified, centered on the somatomotor area. The changes we identified in this network encompasses connections with the amygdala, striatum, and interbrain regions (Figure 6A). Within this network, we identified two centers of plasticity illustrated respectively in panels B and C (Figure 6B-C).

**Figure 6:**
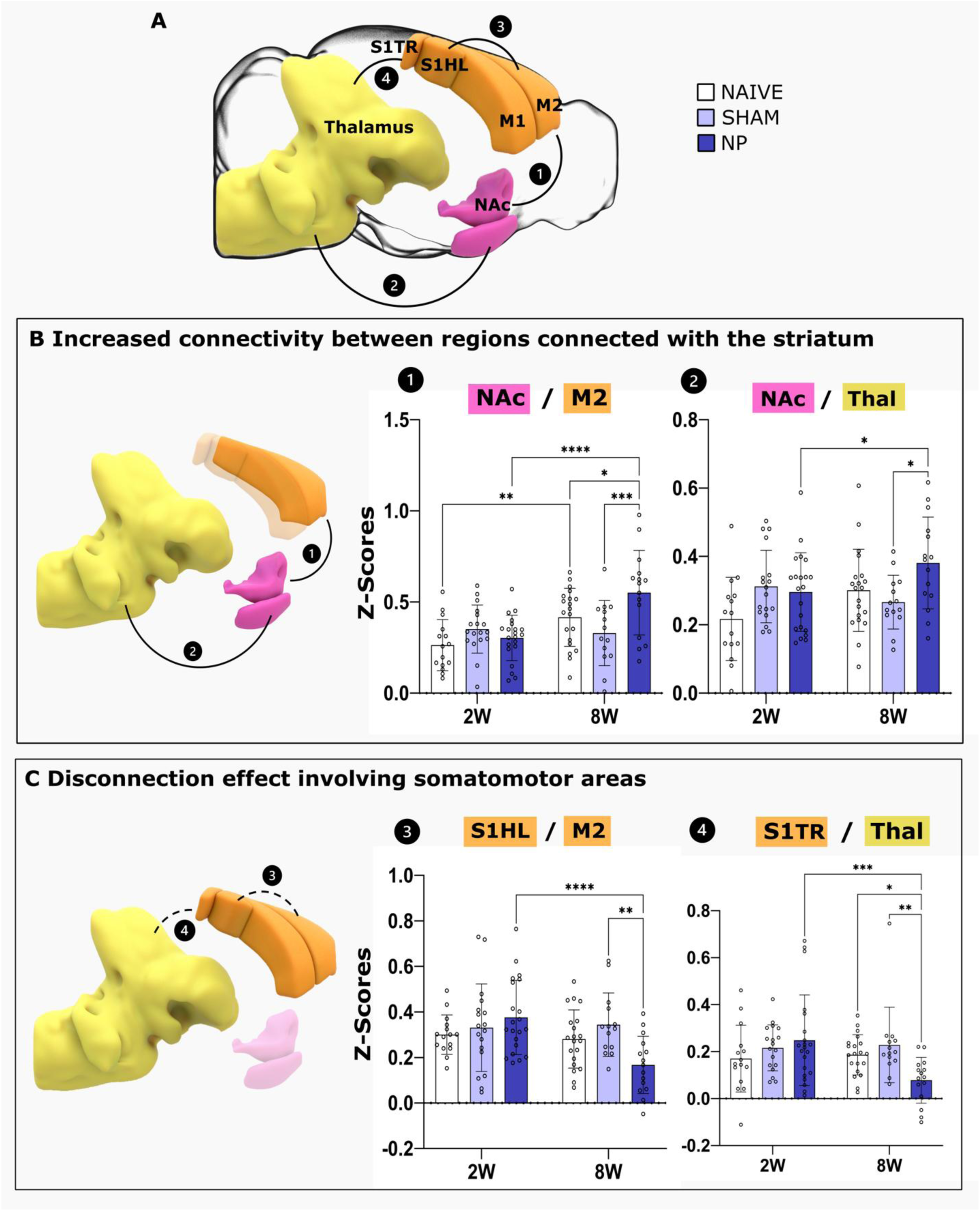
Functional connectivity alterations of a specific subnetwork at 8W. (A) Schematized illustration of the subnetworks showing significant functional connectivity alterations. Somatomotor area, interbrain, amygdala and striatum. (B-C) Division of subnetworks showing an increase in the connectivity between regions connected to the striatum (B) and a disconnection effect involving somatomotor areas (C). The increase in connectivity is illustrated as a bold line and the decrease as a dotted line. The changes illustrated refer to the alterations detected in the neuropathic group compared to the sham group. Graphs 1, 2, 3 and 4 represent Fisher-transformed z-scores of correlations between pairs of ROIs at 2 and 8 weeks. Data are presented as mean ± SD (*p<0.05, **p<0.01, ***p<0.001, ****p<0.0001). Mice n=12, acquisitions n=49. 8W: Naïve n=5, Sham n=5, NP n=2.

First, we identified changes in connectivity strength among regions connected with the striatum (Figure 6B). The neuropathic group shows an increased FC between NAc and M2 (Figure 6B graph 1, p= 0.0002, q=6.092, DF=57 vs SHAM, p= 0.0319, q=3.667, DF=57 vs NAÏVE), and between NAc and thalamus (Figure 6B graph 2, p=0.0263, q=3.775, DF=57 vs SHAM) and to the NP group at 2W (p<0.0001, q=6.476, DF=37 and p=0.0257, q=3.287, DF=37 respectively).

Second, we centered our analysis on the FC from somatomotor areas and functionally related areas and found in contrary a decreased FC in neuropathic animals at 8W in the following pairs of regions: 1) S1HL and M2 (p=0.0033, q=4.839, DF=57; Figure 6C graph 3) and 2) S1TR-Thal (p=0.0022, q=5.021, DF=57 vs sham, p=0.0292, q=3.718, DF=57 vs naïve, Figure 6C graph 4). These variations were also significantly different from the neuropathic group imaged at 2W (NP 2W vs 8W: S1HL-M2 p<0.0001, q=6.345, DF=37 S1TR-Thal p= 0.0004, q=5.497, DF=37). These results show a disconnection within the somatomotor areas, and between the thalamus due to neuropathic pain 8 weeks after the cuff surgery.

These results highlight an opposite plasticity effect concerning these two subnetworks. The neuropathic lesion is associated with a disconnection in the regions linked with the somatomotor cortex and a stronger connection between regions linked with the striatum, 8 weeks after the cuff surgery.

## DISCUSSION

Imaging techniques are instrumental in uncovering the complex neurobiology underlying chronic pain conditions, such as neuropathic pain ^25^. Using functional ultrasound (fUS) imaging in both awake and anesthetized neuropathic mice and their controls, we found that while most brain networks maintain stable functional connectivity (FC), neuropathic pain induces selective, time-specific changes—particularly involving the prefrontal cortex and hypothalamus. Additionally, the persistence of neuropathic pain led to reduced FC within the somatomotor network.

### Stability of Functional Connectivity Across Brain Regions

Our findings demonstrate a notable stability in FC across most brain areas over time. We initially hypothesized that anesthesia might obscure important changes, as suggested in prior studies ^26–28^. However, awake recordings confirmed that only specific networks exhibit FC alterations in neuropathic pain.

Unexpectedly, anesthetized animals showed a transient decrease in FC at two weeks across all groups, followed by increases at 8 and 12 weeks. Since this pattern was consistent across conditions, it is likely due to external experimental factors—most probably the cumulative effects of repeated ketamine administration, known to modulate FC ^29,30^.

### FC Changes Following Surgery and Neuropathic Pain Induction

We observed decreased FC between the infralimbic cortex (IL) and amygdala in both sham and neuropathic groups two weeks post-lesion, relative to naïve mice. Because both groups underwent surgery, this suggests an effect of post-surgical pain. Notably, by 8 and 12 weeks, FC between IL and amygdala returned to baseline in all groups, except neuropathic mice, implying a persistent pain-related mechanism.

The amygdala is central to the emotional aspects of pain—fear, anxiety, and stress responses ^31,32^. The IL, on the other hand, is implicated in cognitive processes such as attentional control, decision-making, and extinction learning, which can modulate the experience of pain. In chronic pain, medial prefrontal cortex (mPFC) activity is typically reduced due to decreased excitatory input and increased GABAergic inhibition ^33^. Disruption of mPFC–amygdala interactions shifts the control of stress responses from top-down to bottom-up mechanisms ^34^.

Clinical studies in pediatric CRPS patients have shown enhanced FC between the amygdala and multiple brain areas ^35^, and similar alterations have been noted in animal models of stress-induced pain ^36^. We propose that the specific biphasic IL–amygdala FC pattern reflects neuroadaptive processes during the transition from maintenance to chronification phases, potentially contributing to comorbidities like anxiety and depression ^16,37^.

### FC Alterations Involving the Insula

The early phase of neuropathic pain was also marked by increased FC between the insula and prelimbic cortex, and decreased FC between the dorsal and posterior insula. The insula, a hub for integrating sensory, emotional, and cognitive information, plays a crucial role in pain processing and interoception ^38^. It is activated in response to noxious stimuli in an intensity-dependent, somatotopically organized manner ^39–41^. Its involvement in aversive behavior, anxiety, and emotional regulation has been extensively described ^42,43^.

Preclinical evidence on the posterior insula’s specific role remains sparse ^44^, though it has been implicated in mechanical allodynia ^45^ and pain aversion ^46^. Our results suggest a functional reorganization of the insular cortex in chronic pain. Increased FC between the insula and prelimbic cortex may represent heightened integration of sensory-affective signals, while decreased intra-insular FC may reflect impaired sensory-affective integration. Together, these changes support the concept of large-scale brain network reorganization in chronic pain.

### Somatomotor Network Dissociation in Chronic Pain

In the chronic phase, we observed FC reductions within the somatomotor network, particularly between S1HL/M2 and S1Tr/thalamus. Similar disconnections have been documented in other pain models and in clinical settings. In a previous study using fUS imaging in anesthetized rats, we identified alterations in the somatomotor cortex in a model of inflammatory pain, with FC changes correlating with pain behaviors and physiological impact ^12^.

Consistent with our findings, altered somatomotor FC has been observed in CRPS and chronic back pain patients, involving thalamic and periaqueductal gray (PAG) connectivity ^47,48^. Interestingly, these changes in our model appeared only at 8 weeks post-lesion, suggesting that long-term neural remodeling in the somatomotor network requires time to manifest. This aligns with reports of delayed electrophysiological alterations in chronic pain models ^49^.

### Increased FC Between NAc and Thalamus During Pain Chronification

The nucleus accumbens (NAc), a central component of the brain’s reward circuitry, comprises a core and shell region ^50^. While the shell is associated with value predictions for monetary gambles, the core is activated during the anticipation of cessation of thermal pain, which is a reward value of pain relief. The NAc is causally involved in the transition to chronic pain in humans ^5,51^ and is necessary for full expression of neuropathic pain-like behavior in rodents ^52^. Notably, only the core NAc has connections to the thalamus in humans, while in rodents, only the shell NAc has strong thalamic connections ^52,53^.

Persistent pain induces significant structural rearrangements of the NAc. In subacute back pain, NAc volume decreases significantly before the development of chronic pain and remains unchanged at follow-up, suggesting that it plays a role in risk for development of chronic pain. In addition, a shift in the electrophysiological properties of the NAc takes place. These alterations in low-frequency (0.01 to 0.027 Hz) oscillations at rest, are a signature of the state of chronic pain in this condition ^54^.

Increased excitatory neurotransmitters content in the thalamus in neuropathic pain conditions contributes to both sensitization and mechanical allodynia ^55^. Moreover, alteration of the FC between the right core NAc and left thalamus was evidenced in patients with fibromyalgia, with a reduced FC in these conditions, in which the reduced reward towards pain relief is well established ^56^. In our study, a reinforcement of the FC between the NAc (core and shell parts) and the thalamus was observed specifically in neuropathic animals. Both these regions are involved in emotional processing and regulation. The dysregulated connectivity observed between these regions may contribute to emotional disturbances and mood disorders associated with neuropathic pain.

### Limitations of our study

Imaging studies in human patients were the first approach employed to uncover dysfunctional brain reorganization. These studies have been critical for identifying the brain circuitry involved in pain processing and modulation, and for understanding the disruption of those circuits in chronic pain. Despite the important information provided by imaging human subjects, there are many limitations. For example, longitudinal or lifespan studies in controlled conditions in humans are very difficult to perform. Conversely, in animal models, especially in rodent imaging studies, it is possible to overcome many of these limitations and provide a mechanism for back-translation of findings from humans to rodent models ^25,57^. In animal models more detailed, controlled, and invasive analysis can be performed. Therefore, we chose to conduct this study in a mouse model of neuropathic pain and longitudinally follow key stages of symptoms development and comorbidity progression within the disease.

To mitigate the bias introduced by anesthesia, we conducted a second set of experiments under awake conditions, taking advantage of fUS imaging’s compatibility with both freely moving and head-fixed animals ^17^. However, this approach also presented challenges. Motion artifacts in awake experiments led to the exclusion of certain acquisitions, reducing the number of animals available for analysis. Despite an initial cohort of 59 mice, only 24 were ultimately included in the final dataset. Furthermore, conducting longitudinal studies in awake conditions proved challenging due to difficulties in maintaining clear cranial windows, necessitating the use of separate cohorts for each time point.

To conclude, this study reveals changes in functional connectivity between key brain regions implicated in pain processing and emotional regulation, namely the infralimbic area and the amygdala, as well as the prelimbic area and the insula. These alterations suggest adaptive changes in brain networks during the chronification of pain and the development of comorbidities, shedding light on the underlying mechanisms involved. These findings offer valuable implications for understanding the neurobiological basis of chronic pain and for the development of targeted therapeutic interventions aimed at alleviating both the physical and emotional burden experienced by chronic pain patients.

## Supporting information

The text is in the PDF

## Authors contribution statement

SP, JF and IY designed the experimental paradigm.

SC performed the functional ultrasound imaging in both awake and anesthetized animals.

SC, JF and SP wrote the manuscript.

MT, TD and BFO supervised the signal processing of the ultrasound data.

SLMED, AB, LE and SC performed the signal processing of the ultrasound data.

SP, SC and SLMED performed the statistical analysis.

SC, JF, IY and SP were involved in the interpretation of the data and wrote different parts of the manuscript.

## Acknowledgments

This work was supported by Iconeus (S. Cazzanelli’s PhD fellowship), the ANRT (S. Cazzanelli’s PhD fellowship), the Agence Nationale de la recherche (Project ‘PINCH’, 18-CE37-0005-01), the Institut National de la Santé et de la Recherche Médicale, CNRS, ESPCI-Paris and PSL University.

Authors would like to thank Mrs L. Antoinette for animal husbandry. Finally, this work was supported by the Inserm ART (Technology Research Accelerator) in "Biomedical Ultrasound".

Several panels of figures were made using Biorender (Biorender.com).

## Data availability statement

Source data are available in the Zenodo repository at the following link: 10.5281/zenodo.15837352.

This dataset includes all individual values corresponding to the figures, as well as the time series of functional connectivity (FC) changes in naïve animals (both anaesthetised and awake). Due to file number limitations on the repository, the time series for injured animals could not be uploaded. However, these data are available upon request from the corresponding author.

## Code availability statement

This work did not involve any custom code. Data acquisition and processing was performed using commercially available softwares (Icoscan, Icostudio, Icolab, Excel, Prism).

---

**Supplementary Figure 1:**
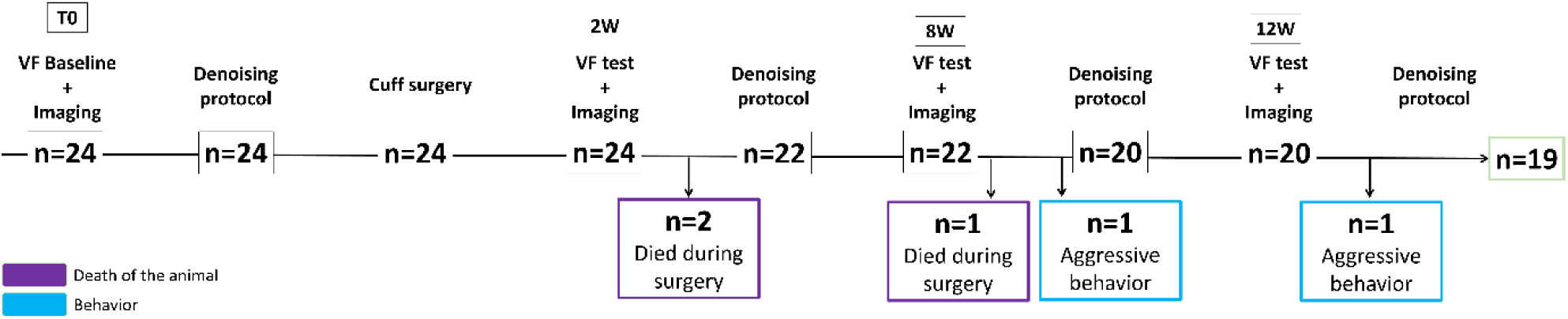
Anesthetized project timeline illustrated as a Gantt Chart indicating stages where animals were excluded for specific reasons. The horizontal axis displays time in weeks, along with the number of mice remaining in the study at each designated time point (T0, 2W, 8W, 12W). The study initially included n=24 mice. At 2 weeks n=2 mice were excluded from the study because they died during surgery. At 8 weeks, n=1 mouse died during surgery, and n=1 was excluded due to aggressive behavior. At 12 weeks, n=1 mouse was excluded for similar behavioral issues. The study concluded with n=19 mice.

**Supplementary Figure 2:**
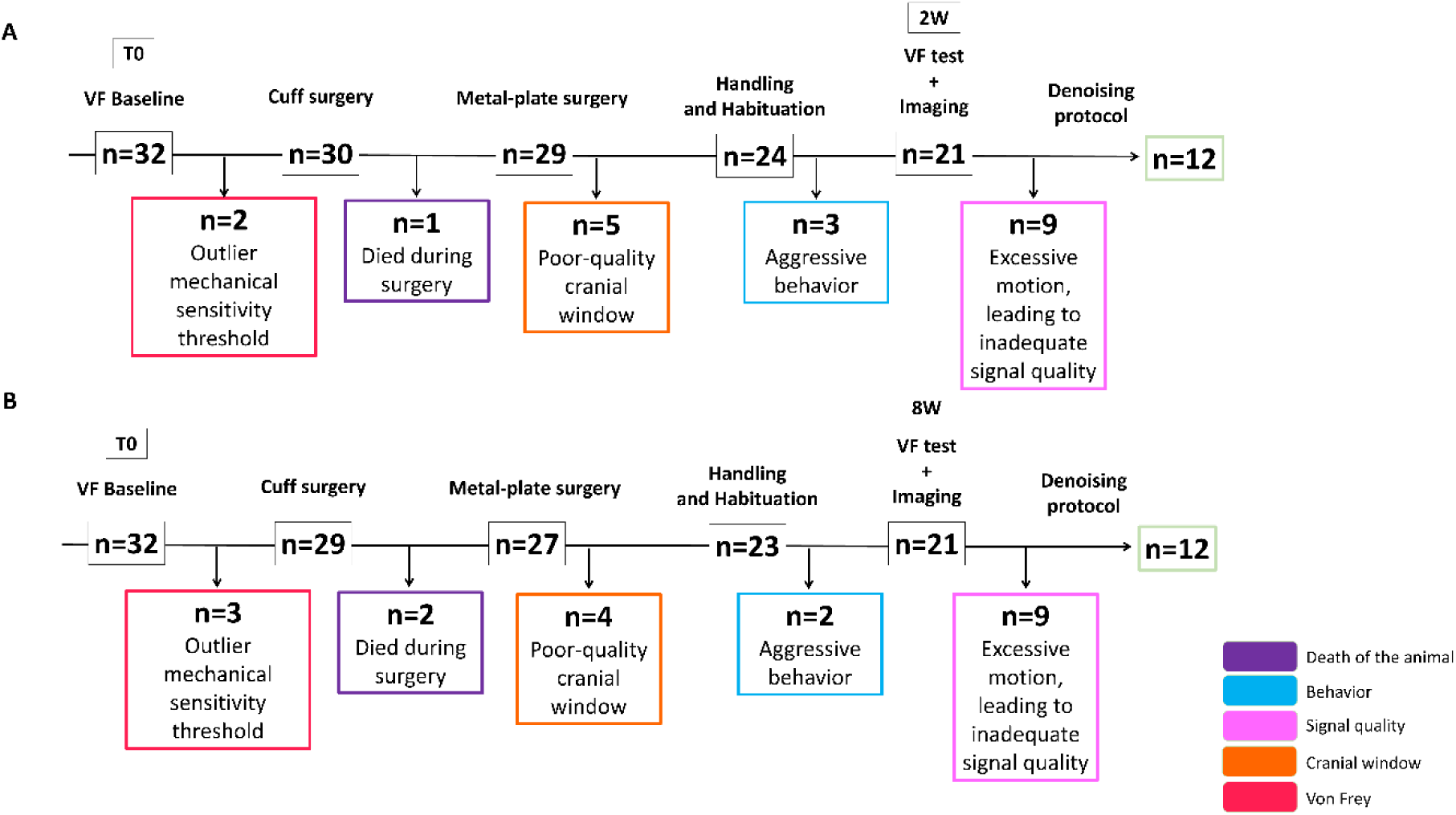
Awake project timelines illustrated in a Gantt Chart indicating stages where animals were excluded for specific reasons. Schematic illustration of two timelines corresponding to two timepoints investigated separately (A-B). The horizontal axis displays time in weeks, along with the number of mice remaining in the study. (A) The 2 weeks study initially included n=32 mice. N=2 were excluded from the study because of outlier mechanical sensitivity threshold. N=1 mice were excluded from the study because they died during surgery. N=5 were excluded from the study because of cranial window’s poor quality, N=3 were excluded because of aggressive behavior and finally n=9 had to be removed from the study after the denoising protocol because of an excessive motion, which leads to a noisy and inadequate signal quality. The study concluded with n=12 mice. (B) The 8 weeks study initially included n=32 mice. N=3 were excluded from the study because of outlier mechanical sensitivity threshold. N=2 mice were excluded from the study because they died during surgery. N=4 were excluded from the study because of cranial window’s poor quality, N=2 were excluded because of aggressive behavioral and finally n=9 had to be removed from the study after the denoising protocol because of an excessive motion, which leads to a noisy and inadequate signal quality. The study concluded with n=12 mice.

**Supplementary Figure 3:**
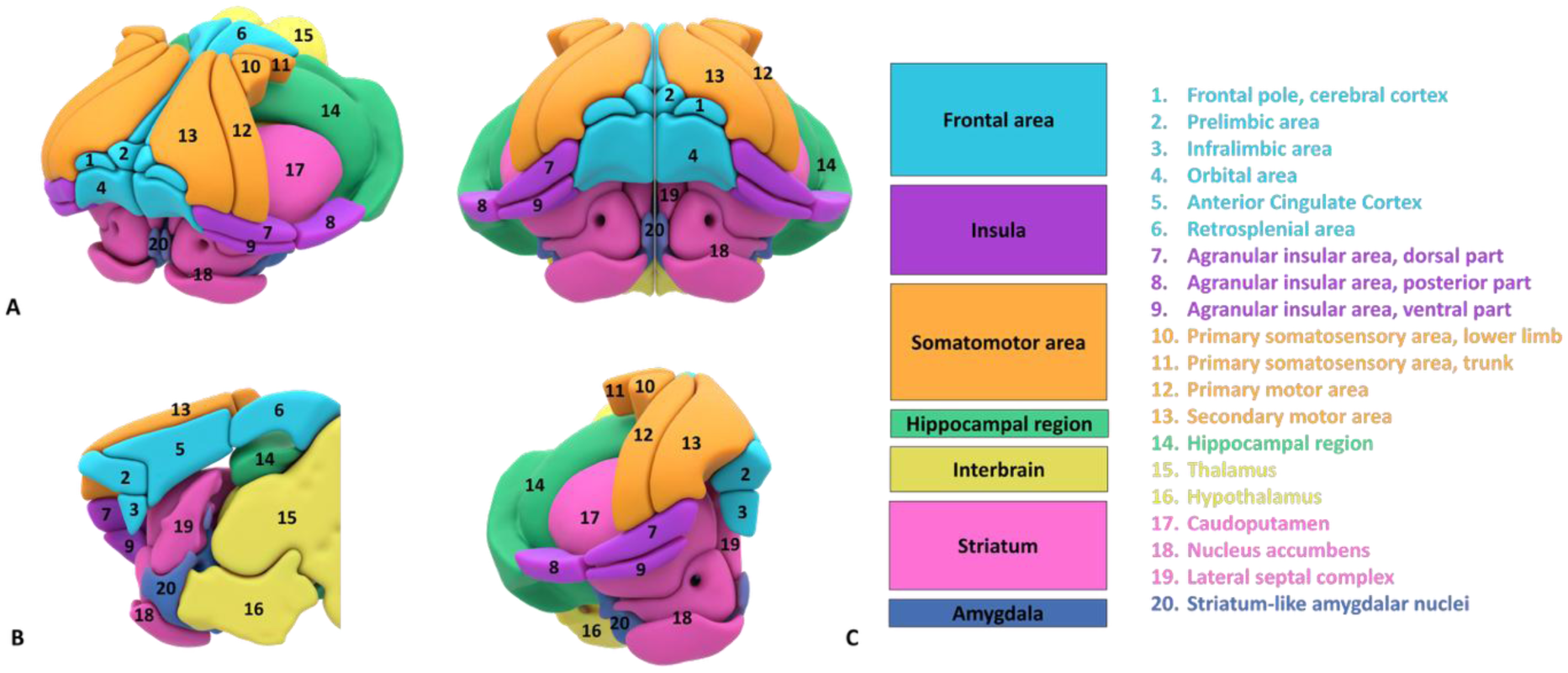
Schematic of the brain areas studied in anesthetized (A) and awake. (B) experimental conditions. C: List of the 20 regions of interest, grouped into 7 macro-regions. In the frontal area are grouped the frontal pole, prelimbic, infralimbic and orbital area, the anterior cingulate cortex and the retrosplenial cortex. The macro-region referred to as the insula includes the dorsal, posterior, and ventral parts of this brain region. The somatomotor area includes the primary somatosensory area corresponding to the lower limb and trunk, as well as the primary and secondary motor areas. The macro-region referred to as interbrain comprises the thalamus and the hypothalamus. The striatum includes the caudoputamen, nucleus accumbens and the lateral septal complex.

**Supplementary Figure 4:**
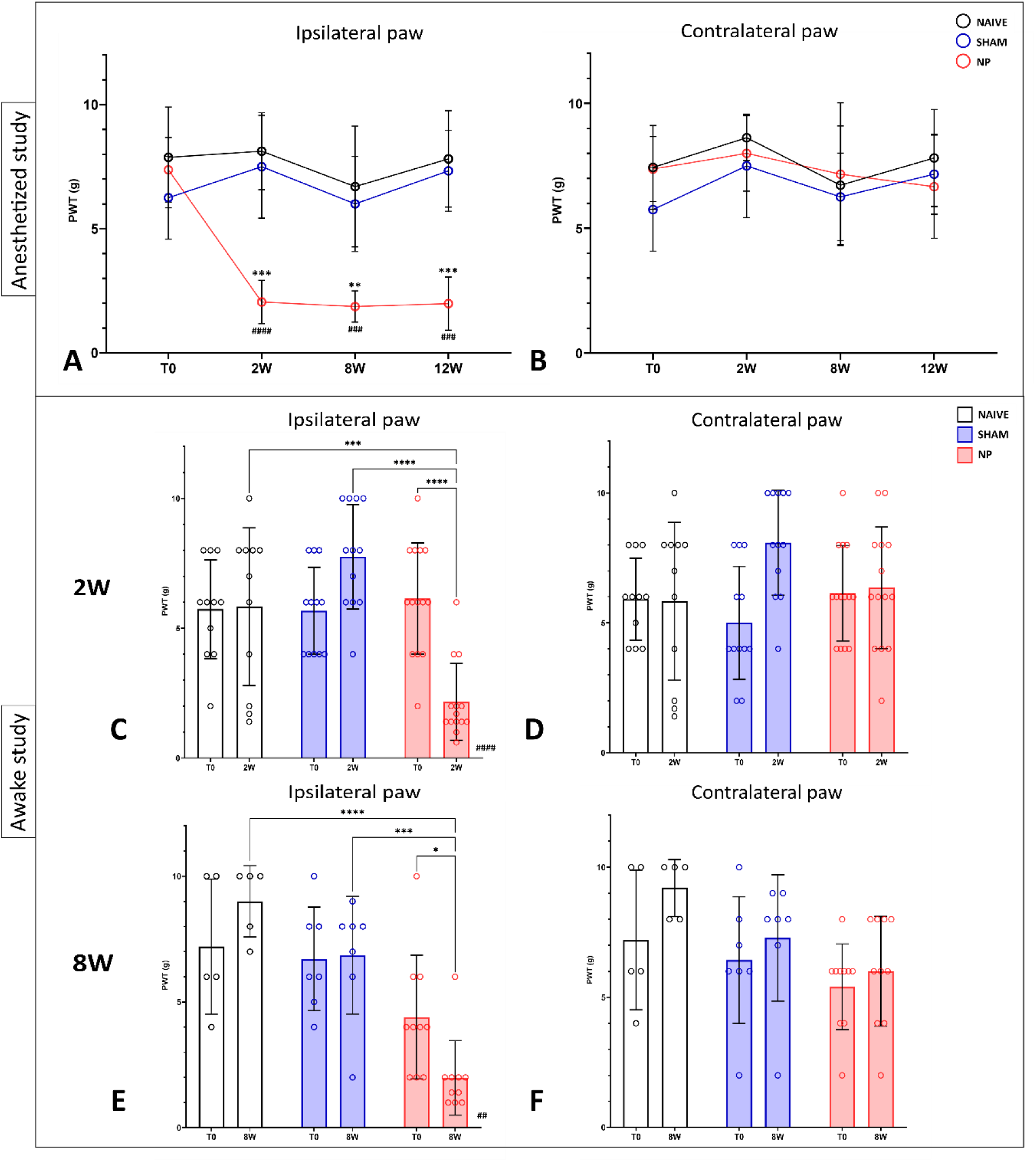
Cuff implantation induced mechanical allodynia. (A-B) Development of mechanical allodynia during the anesthetized study. The Von Frey test was carried out at each time point on the ipsilateral (A) and contralateral (B) hind paw showing the development of mechanical allodynia in the neuropathic group’s ipsilateral paw starting at 2W and its persistence at 8W and 12W. Data are expressed as mean ± SD (* comparison between NP and Sham ipsilateral paws, ** p<0.01, *** p<0.001) (# comparison with NP at T0, 11### p<0.001, #### p<0.0001) (T0: Naïve n=8, Sham n=8, NP n=8; 2W: Naïve n=8, Sham n=7, NP n=8; 8W: Naïve n=8, Sham n=7, NP n=6; 12W: Naïve n=7, Sham n=6, NP n=6). (C-F) Development of mechanical allodynia during the awake study. This part of the study required two batches of animals, Batch #1 (panels C-D): Naïve n=11, Sham n=12, NP n=14 and Batch #2 (panels E-F) Naïve n=5, Sham n=7, NP n=10. For both batches the mechanical sensitivity was tested on the ipsilateral (C and E) and the contralateral paw (D and F) at baseline (T0) and at the corresponding timepoints: for Batch #1 at 2W (C-D) and for Batch #2 at 8W (E-F). Data are expressed as mean ± SD (* comparison between Naïve vs NP and Sham vs NP ipsilateral paws, *** p<0.001, ****p<0.0001) (# comparison between T0 and 2W or 8W NP group ipsilateral paws. #p<0.005, #### p<0.0001) ($ comparison between ipsilateral and contralateral paws of NP group at 2W or 8W, $$p<0.01 $$$$p<0.0001).

**Supplementary figure 5:**
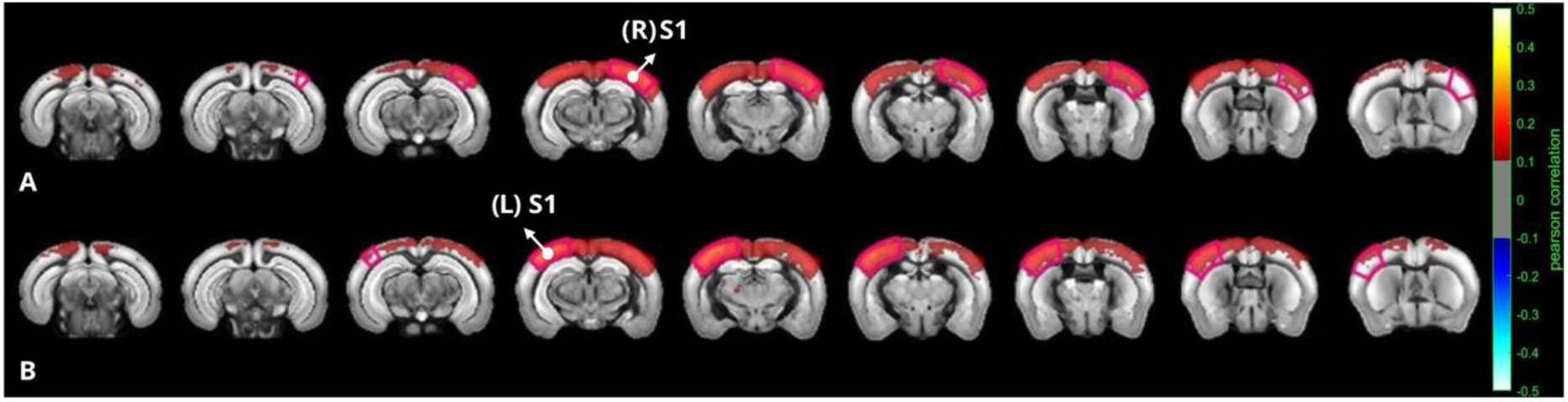
rsFC network using seed-based analysis. Group-averaged seed-based correlation maps in anesthetized mice. The two seeds are the right part of the primary sensory cortex (R-S1) (A) and its bilateral part (L-S1) (A). Correlation map is obtained by computing the Pearson Correlation coefficient between the temporal signals of the seed and each voxel of the whole acquisition after slice timing correction.

**Supplementary figure 6:**
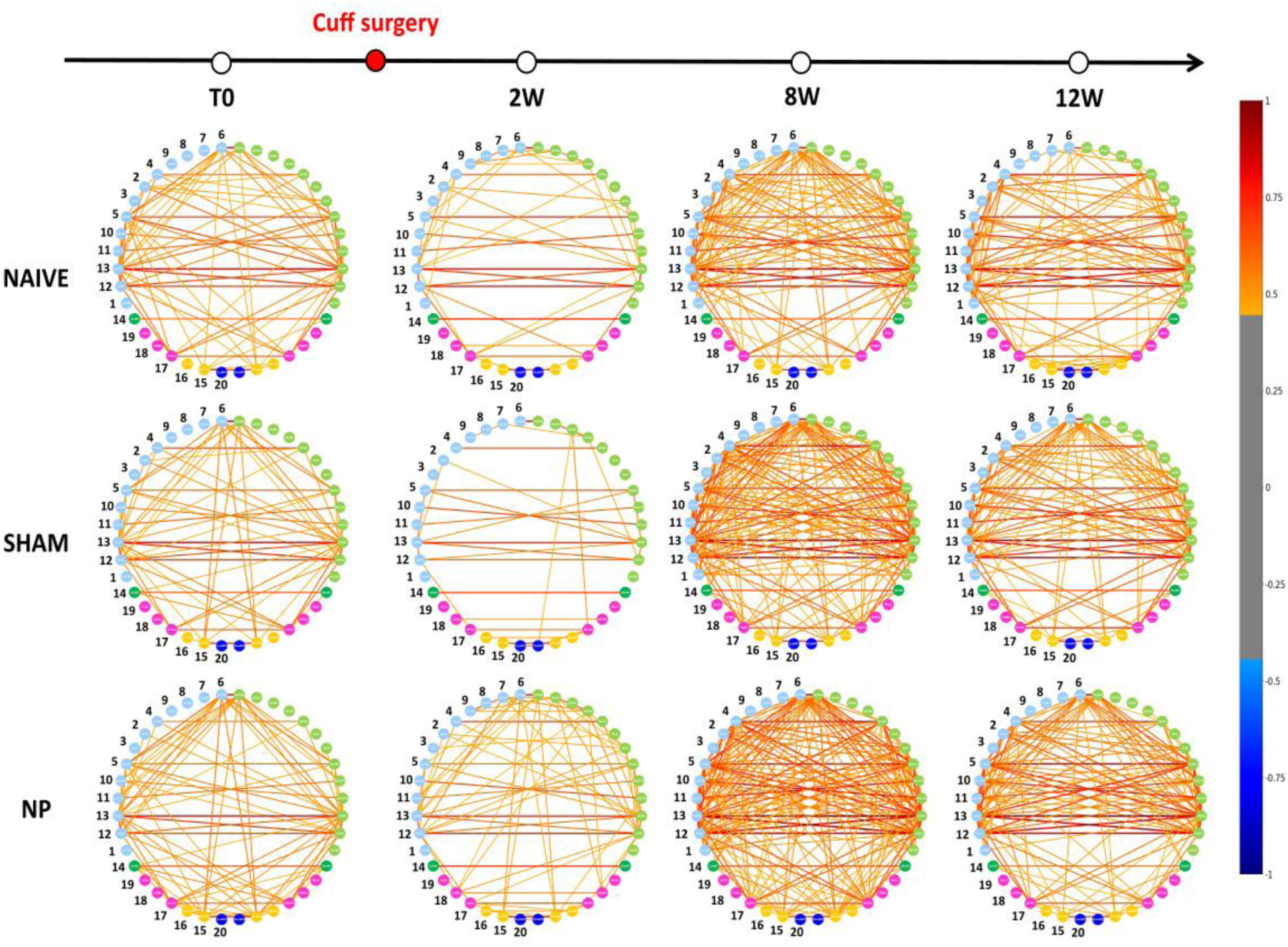
Circular networks showing Functional Connectivity alterations in a wide-range network. Circular network representation. Alternative representation of the correlation matrix as a circular network with the 40 ROIs displayed in a circular layout and the connection between them illustrated as links which color corresponds to the correlation coefficient between the two nodes. The 40 ROIs are listed in Table 1A. T0: Naïve n=8, Sham n=8, NP n=8; 2W: Naïve n=7, Sham n=7, NP n=8; 8W: Naïve n=8, Sham n=6, NP n=6; 12W: Naïve n=7, Sham n=6, NP n=6.

**Supplementary Figure 7:**
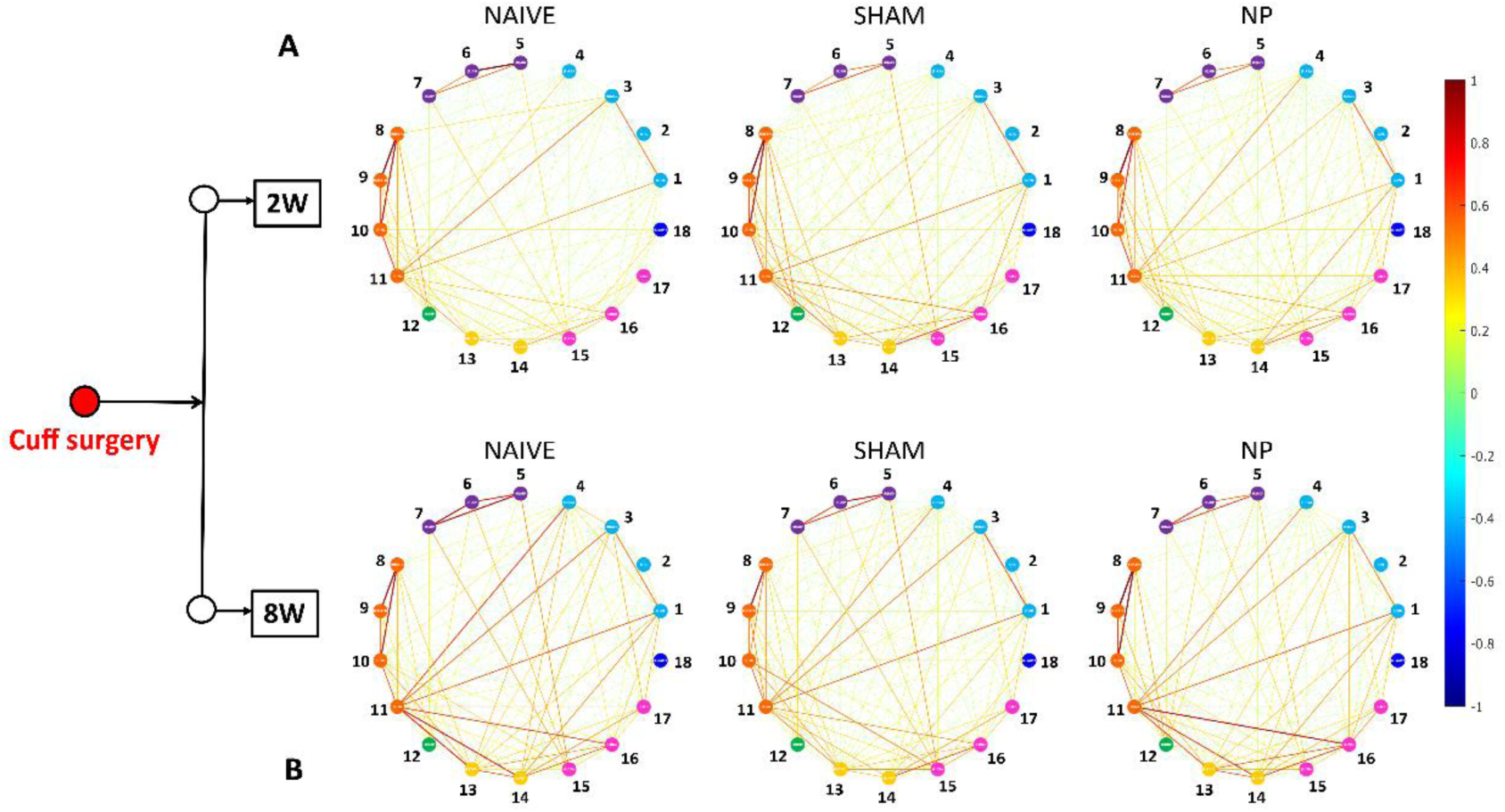
Functional Connectivity alterations in a wide-range network. (A-B-D) Alternative representation of the correlation matrix as a circular network with the 18 ROIs displayed in a circular layout and the connection between them illustrated as links color corresponds to the correlation coefficient between the two nodes. The 18 ROIs are listed in Table 1B. 2W: Naïve n=2, Sham n=5, NP n=5; 8W: Naïve n=5, Sham n=5, NP n=2.

