## Supplementary material for "Temporal Patterns of Brain Network Plasticity During the Onset and Maintenance of Neuropathic Pain in Male Mice": The text is in the PDF

**Cazzanelli et al., SUPPLEMENTARY MATERIALS**

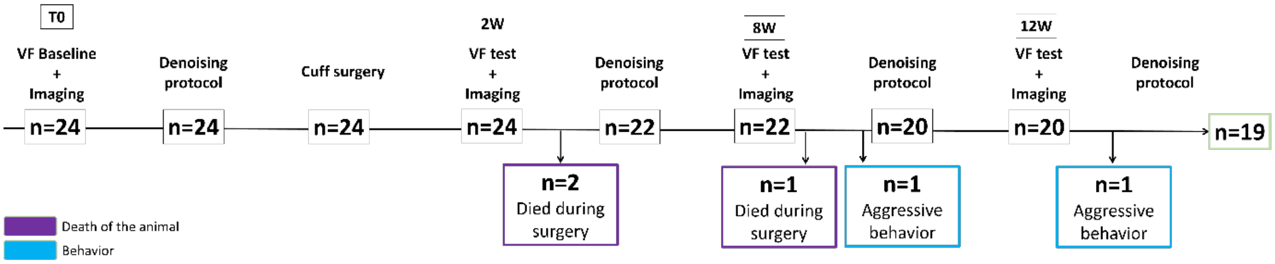


**Supplementary Figure 1: Anesthetized project timeline illustrated as a Gantt Chart indicating stages where animals were excluded for specific reasons.** The horizontal axis displays time in weeks, along with the number of mice remaining in the study at each designated time point (T0, 2W, 8W, 12W). The study initially included n=24 mice. At 2 weeks n=2 mice were excluded from the study because they died during surgery. At 8 weeks, n=1 mouse died during surgery, and n=1 was excluded due to aggressive behavior. At 12 weeks, n=1 mouse was excluded for similar behavioral issues. The study concluded with n=19 mice.

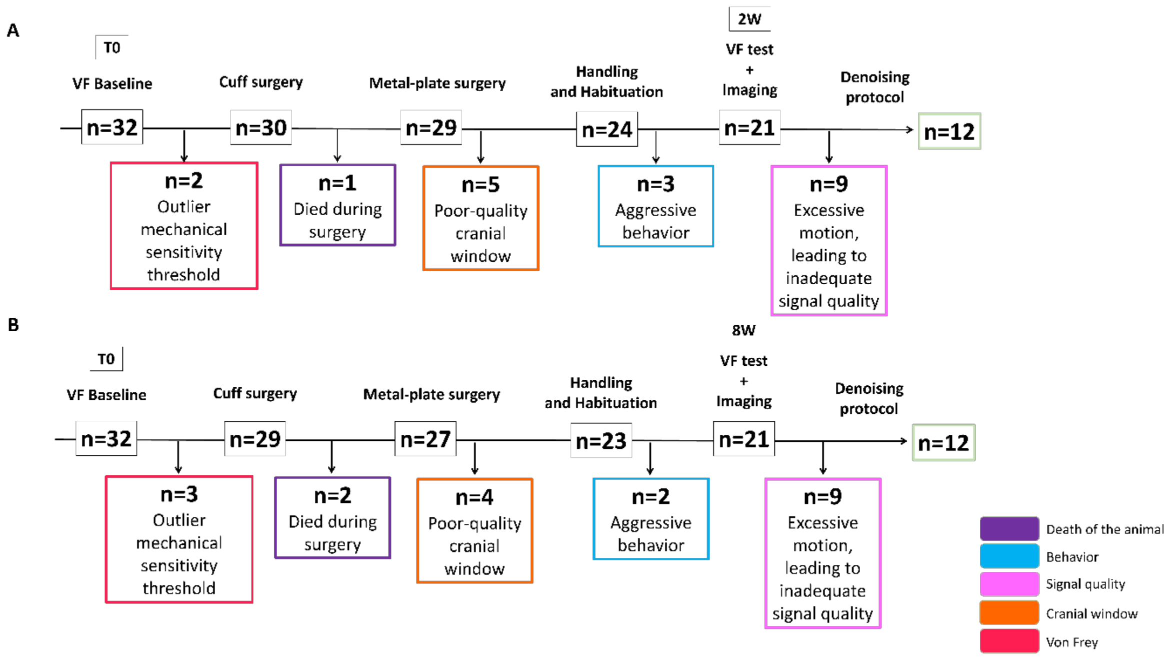


**Supplementary Figure 2: Awake project timelines illustrated in a Gantt Chart indicating stages where animals were excluded for specific reasons.** Schematic illustration of two timelines corresponding to two timepoints investigated separately (A-B). The horizontal axis displays time in weeks, along with the number of mice remaining in the study. (A) The 2 weeks study initially included n=32 mice. N=2 were excluded from the study because of outlier mechanical sensitivity threshold. N=1 mice were excluded from the study because they died during surgery. N=5 were excluded from the study because of cranial window's poor quality, N=3 were excluded because of aggressive behavior and finally n=9 had to be removed from the study after the denoising protocol because of an excessive motion, which leads to a noisy and inadequate signal quality. The study concluded with n=12 mice. (B) The 8 weeks study initially included n=32 mice. N=3 were excluded from the study because of outlier mechanical sensitivity threshold. N=2 mice were excluded from the study because they died during surgery. N=4 were excluded from the study because of cranial window's poor quality, N=2 were excluded because of aggressive behavioral and finally n=9 had to be removed from the study after the denoising protocol because of an excessive motion, which leads to a noisy and inadequate signal quality. The study concluded with n=12 mice.

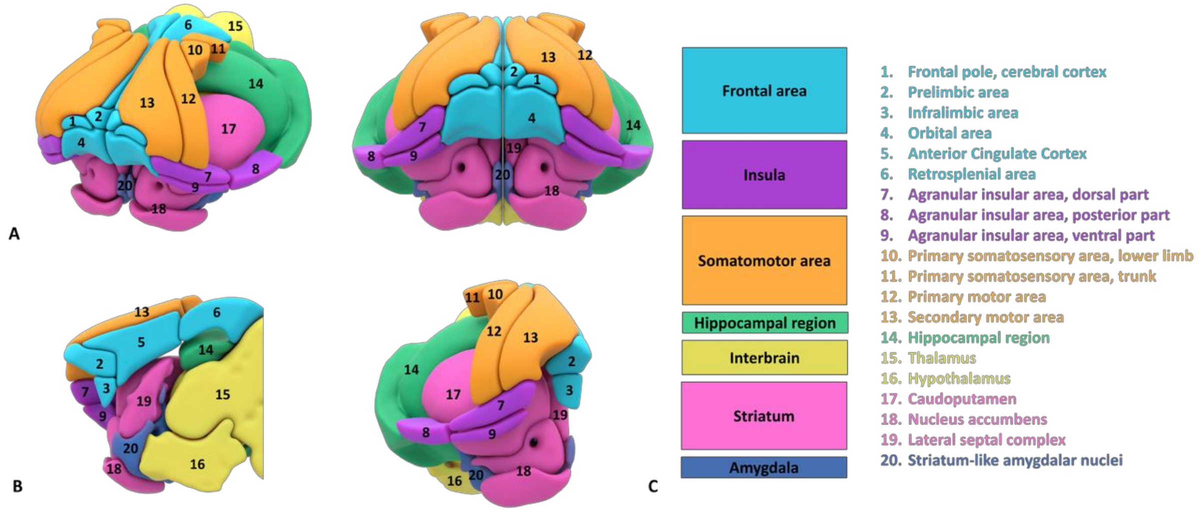


**Supplementary Figure 3: Schematic of the brain areas studied in anesthetized (A) and awake** (B) experimental conditions. C: List of the 20 regions of interest, grouped into 7 macro-regions. In the frontal area are grouped the frontal pole, prelimbic, infralimbic and orbital area, the anterior cingulate cortex and the retrosplenial cortex. The macro-region referred to as the insula includes the dorsal, posterior, and ventral parts of this brain region. The somatomotor area includes the primary somatosensory area corresponding to the lower limb and trunk, as well as the primary and secondary motor areas. The macro-region referred to as interbrain comprises the thalamus and the hypothalamus. The striatum includes the caudoputamen, nucleus accumbens and the lateral septal complex.

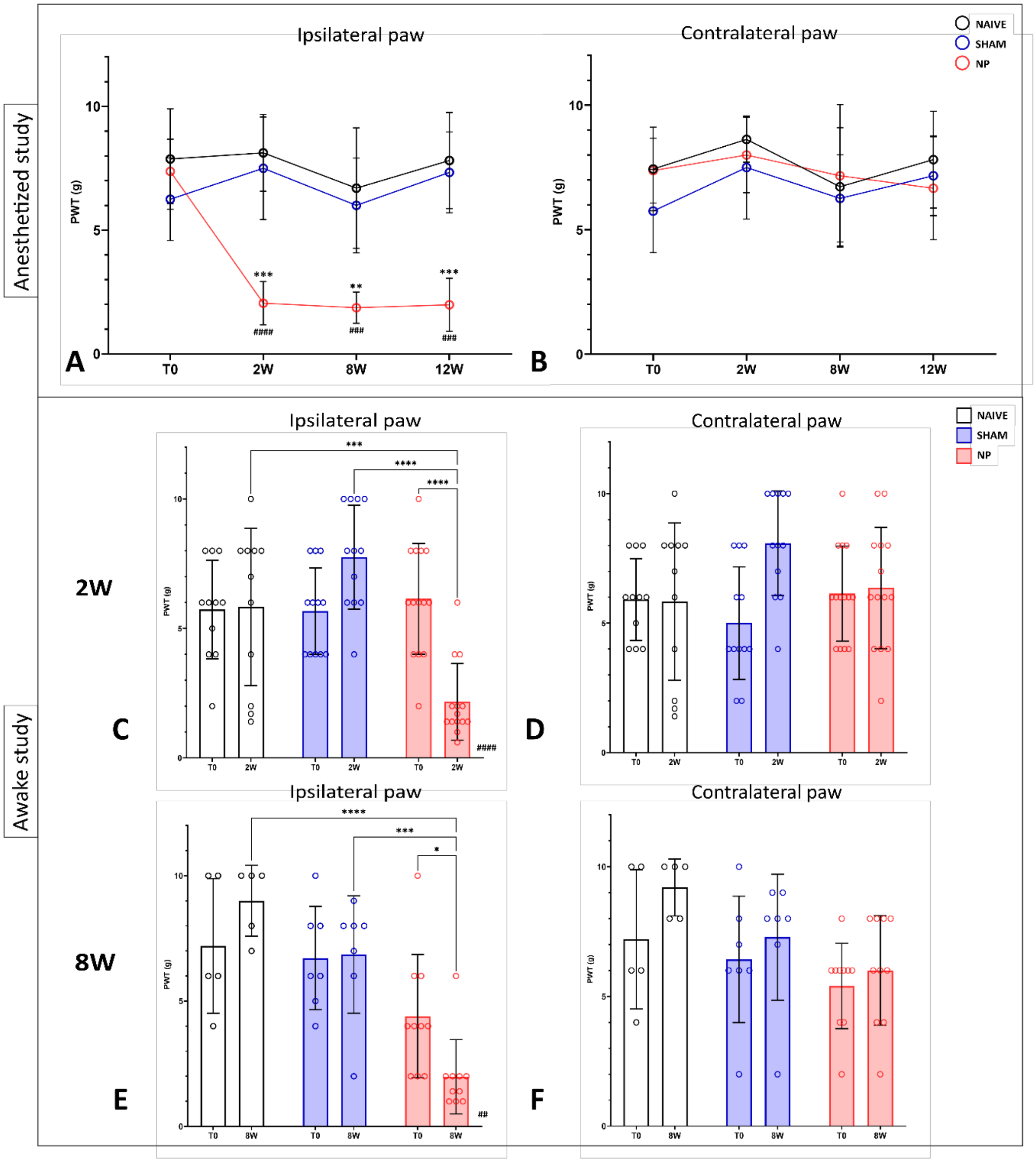


**Supplementary Figure 4: Cuff implantation induced mechanical allodynia.**

(A-B) Development of mechanical allodynia during the anesthetized study. The Von Frey test was carried out at each time point on the ipsilateral (A) and contralateral (B) hind paw showing the development of mechanical allodynia in the neuropathic group's ipsilateral paw starting at 2W and its persistence at 8W and 12W. Data are expressed as mean ± SD (* comparison between

NP and Sham ipsilateral paws, ** p<0.01, *** p<0.001) (# comparison with NP at T0, 11### p<0.001, #### p<0.0001) (T0: Naïve n=8, Sham n=8, NP n=8; 2W: Naïve n=8, Sham n=7, NP n=8; 8W: Naïve n=8, Sham n=7, NP n=6; 12W: Naïve n=7, Sham n=6, NP n=6). (C-F) Development of mechanical allodynia during the awake study. This part of the study required two batches of animals, Batch #1 (panels C-D): Naïve n=11, Sham n=12, NP n=14 and Batch #2 (panels E-F) Naïve n=5, Sham n=7, NP n=10. For both batches the mechanical sensitivity was tested on the ipsilateral (C and E) and the contralateral paw (D and F) at baseline (T0) and at the corresponding timepoints: for Batch #1 at 2W (C-D) and for Batch #2 at 8W (E-F). Data are expressed as mean ± SD (* comparison between Naïve vs NP and Sham vs NP ipsilateral paws, *** p<0.001, ****p<0.0001) (# comparison between T0 and 2W or 8W NP group ipsilateral paws. #p<0.005, #### p<0.0001) ($ comparison between ipsilateral and contralateral paws of NP group at 2W or 8W, $$p<0.01 $$$$p<0.0001).

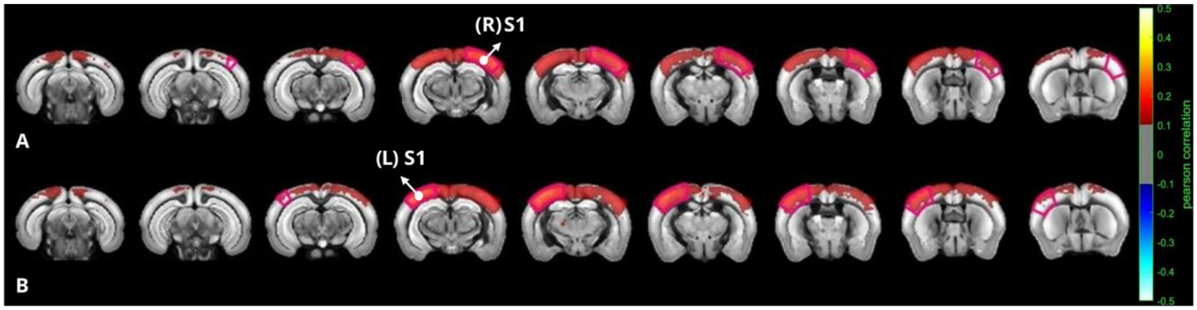


**Supplementary figure 5: rsFC network using seed-based analysis.** Group-averaged seed-based correlation maps in anesthetized mice. The two seeds are the right part of the primary sensory cortex (R-S1) (A) and its bilateral part (L-S1) (A). Correlation map is obtained by computing the Pearson Correlation coefficient between the temporal signals of the seed and each voxel of the whole acquisition after slice timing correction.

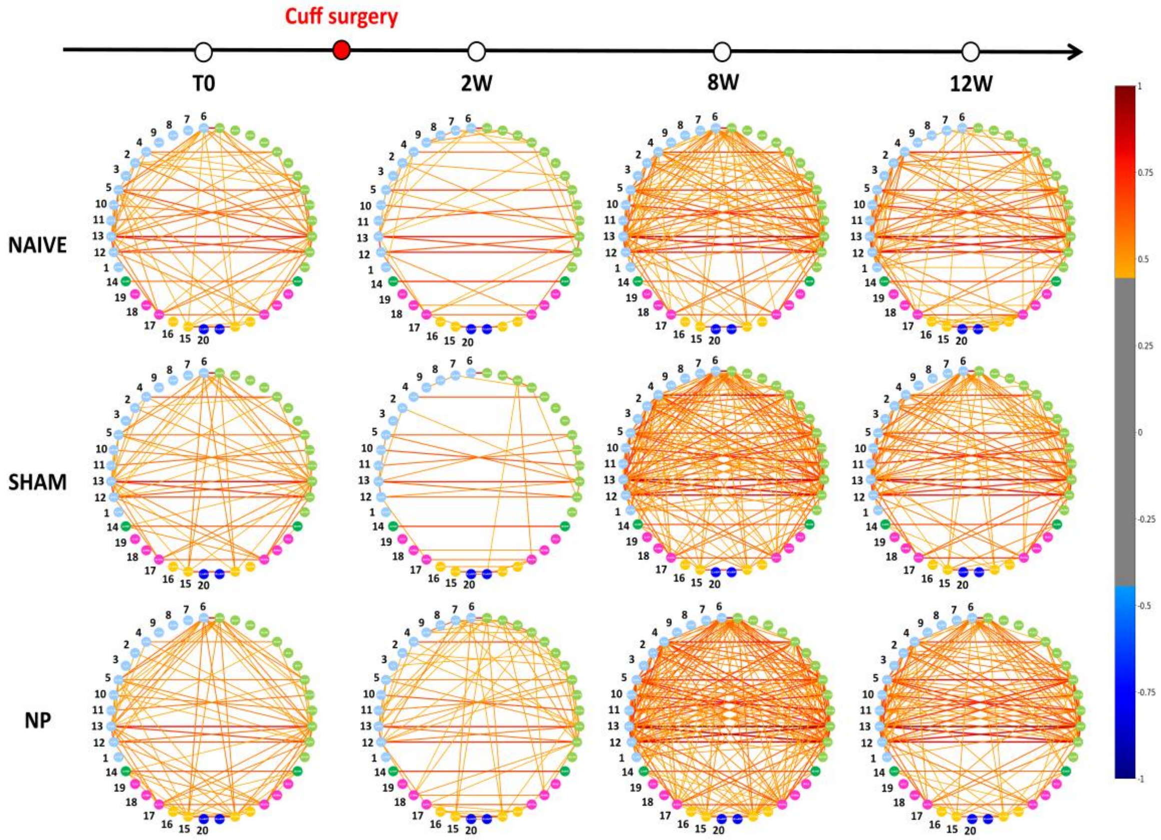


**Supplementary figure 6:** Circular networks showing Functional Connectivity alterations in a wide-range network. Circular network representation. Alternative representation of the correlation matrix as a circular network with the 40 ROIs displayed in a circular layout and the connection between them illustrated as links which color corresponds to the correlation coefficient between the two nodes. The 40 ROIs are listed in Table 1A. T0: Naïve n=8, Sham n=8, NP n=8; 2W: Naïve n=7, Sham n=7, NP n=8; 8W: Naïve n=8, Sham n=6, NP n=6; 12W: Naïve n=7, Sham n=6, NP n=6.

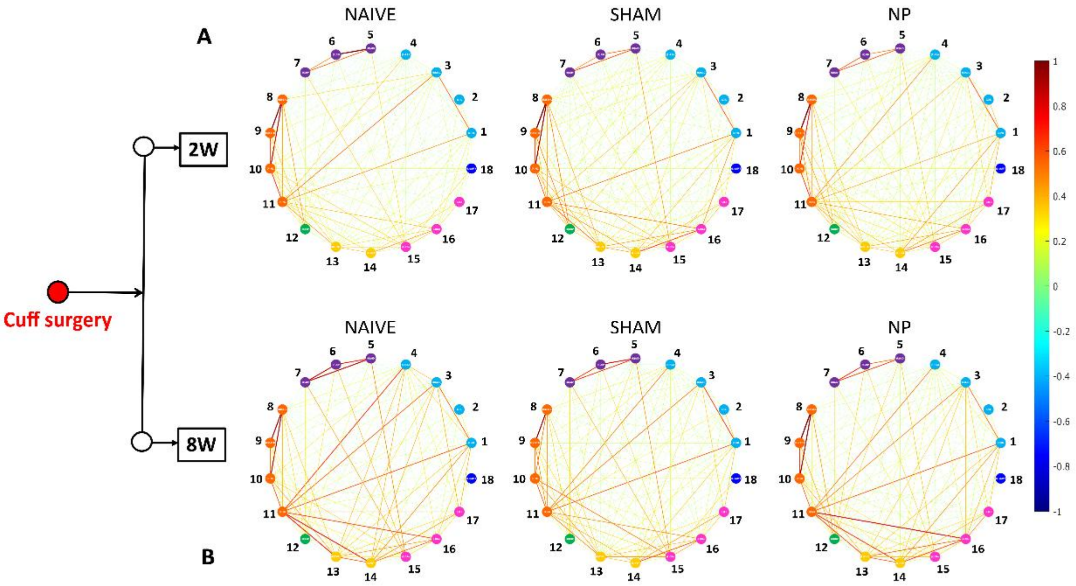


**Supplementary Figure 7:** Functional Connectivity alterations in a wide-range network. (A-B-D) Alternative representation of the correlation matrix as a circular network with the 18 ROIs displayed in a circular layout and the connection between them illustrated as links color corresponds to the correlation coefficient between the two nodes. The 18 ROIs are listed in Table 1B. 2W: Naïve n=2, Sham n=5, NP n=5; 8W: Naïve n=5, Sham n=5, NP n=2.

| Anesthetized configuration | Macro-region | | Bran region | n | Side |
|---|---|---|---|---|---|
| | Isocortex | Frontal area | Frontal pole, cerebral cortex | 1 | right and left |
| | | | Prelimbic area | 2 | right and left |
| | | | Infralimbic area | 3 | right and left |
| | | | Orbital area | 4 | right and left |
| | | | Anterior Cingulate Cortex | 5 | right and left |
| | | | Retrosplenial area | 6 | right and left |
| | | Insula | Agranular insular area, dorsal part | 7 | right and left |
| | | | Agranular insular area, posterior part | 8 | right and left |
| | | | Agranular insular area, ventral part | 9 | right and left |
| | | Somatomotor area | Primary somatosensory area, hind limb | 10 | right and left |
| | | | Primary somatosensory area, trunk | 11 | right and left |
| | | | Primary motor area | 12 | right and left |
| | | | Secondary motor area | 13 | right and left |
| | Hippocampal region | | Hippocampal region | 14 | right and left |
| | Interbrain | | Thalamus | 15 | right and left |
| | | | Hypothalamus | 16 | right and left |
| | Striatum | | Caudoputamen | 17 | right and left |
| | | | Nucleus accumbens | 18 | right and left |
| | | | Lateral septal complex | 19 | right and left |
| | Amygdala | | Striatum-like amygdalar nuclei | 20 | right and left |

A

| Awake configuration | Macro-region | Bran region | n | Side |
|---|---|---|---|---|
| | Frontal area | Prelimbic area | 1 | left |
| | | Infralimbic area | 2 | left |
| | | Anterior Cingulate Cortex | 3 | left |
| | | Retrosplenial area | 4 | left |
| | Insula | Agranular insular area, dorsal part | 5 | left |
| | | Agranular insular area, posterior part | 6 | left |
| | | Agranular insular area, ventral part | 7 | left |
| | Somatomotor area | Primary somatosensory area, hind limb | 8 | left |
| | | Primary somatosensory area, trunk | 9 | left |
| | | Primary motor area | 10 | left |
| | | Secondary motor area | 11 | left |
| | Hippocampal region | Hippocampal region | 12 | left |
| | Interbrain | Thalamus | 13 | left |
| | | Hypothalamus | 14 | left |
| | Striatum | Caudoputamen | 15 | left |
| | | Nucleus accumbens | 16 | left |
| | | Lateral septal complex | 17 | left |
| | Amygdala | Striatum-like amygdalar nuclei | 18 | left |

B

**Table 1: List of Regions of Interest (ROIs).** (A) For the anesthetized configuration, the 40 ROIs are grouped into five macro-regions: the Isocortex, which includes the frontal area, insula, and somatomotor area; the hippocampal region; the interbrain; the striatum; and the amygdala. In this configuration, the brain was imaged bilaterally, with each ROI divided into right and left hemispheric parts. (B) For the awake configuration, the 18 ROIs are grouped into seven macro-regions: the frontal area, insula, somatomotor area, the hippocampal region, the interbrain, the striatum, and the amygdala. In this configuration, the brain was imaged on its left hemisphere.
